# Deep-learning triage of 3D pathology datasets for comprehensive and efficient pathologist assessments

**DOI:** 10.1101/2025.07.20.665804

**Authors:** Gan Gao, Renao Yan, Andrew H. Song, Huai-Ching Hsieh, Lindsey A. Erion Barner, Fiona Wang, David Brenes, Sarah S.L. Chow, Rui Wang, Kevin W. Bishop, Yongjun Liu, Xavier Farre, Mukul Divatia, Michelle R. Downes, Funda Vakar-Lopez, Priti Lal, Wynn Burke, Anant Madabhushi, Lawrence D. True, Deepti M. Reddi, William M. Grady, Faisal Mahmood, Jonathan T.C. Liu

## Abstract

Standard-of-care slide-based 2D histopathology severely undersamples spatially heterogeneous tissue specimens, with each thin 2D section representing <1% of the entire tissue volume (in the case of a biopsy). Recent advances in non-destructive 3D pathology, such as open-top light-sheet microscopy (OTLS), enable comprehensive high-resolution imaging of large clinical specimens. While fully automated computational analyses of such 3D pathology datasets are being explored, a potential low-risk route for accelerated clinical adoption would be to continue to rely upon pathologists to provide final diagnoses. Since manual review of these massive and complex 3D datasets is infeasible for routine clinical practice, we present CARP3D, a deep learning triage framework that identifies high-risk 2D cross sections within large 3D pathology datasets to enable time-efficient pathologist evaluation. CARP3D assigns risk scores to all 2D levels within a tissue volume by leveraging context from a subset of neighboring depth levels, outperforming models in which predictions are based on isolated 2D levels. In two use cases – risk stratification based on prostate cancer biopsies and screening for dysplasia/cancer in endoscopic biopsies of Barrett’s esophagus – AI-triaged 3D pathology, enabled by CARP3D, demonstrates the potential to improve the detection of high-risk diseases in comparison to slide-based 2D histopathology while optimizing pathologist workloads.

## Introduction

The diagnostic standard of care for most medical conditions involves histopathology, where a limited number of thin (2D) tissue sections (typically < 5 *µm* in thickness) are cut from an arbitrary side of a tissue specimen (usually a few *mm* in thickness) and mounted on glass slides for microscopic evaluation. Due to the severe under-sampling of tissue volumes visualized by standard histopathology, and the absence of 3D morphological context, the resulting diagnoses can be inaccurate or ambiguous ^1, 2^ (**Fig. 1a**). For example, studies have revealed that conventional 2D histology sections may fail to capture or accurately represent diagnostically important tissue morphologies, which are often spatially heterogeneous ^3–12^. Recent advances in high-throughput and non-destructive three-dimensional (3D) microscopy ^13–20^, along with rapid and gentle tissue-clearing and fluorescence-labeling techniques ^13, 21–25^, enable comprehensive sampling of large tissue volumes, including whole biopsies. With appropriate stains and robust image-processing methods ^26, 27^, high-quality 3D pathology datasets can be generated that mimic the appearance of hematoxylin and eosin (H&E)-stained tissue sections. This provides orders-of-magnitude more data than 2D histopathology along with valuable 3D information that does not exist within 2D slides ^12, 28, 29^. However, manual review of such large 3D pathology datasets (equivalent to hundreds to thousands of 2D sections per specimen) is impractical in clinical settings.

**Figure 1:**
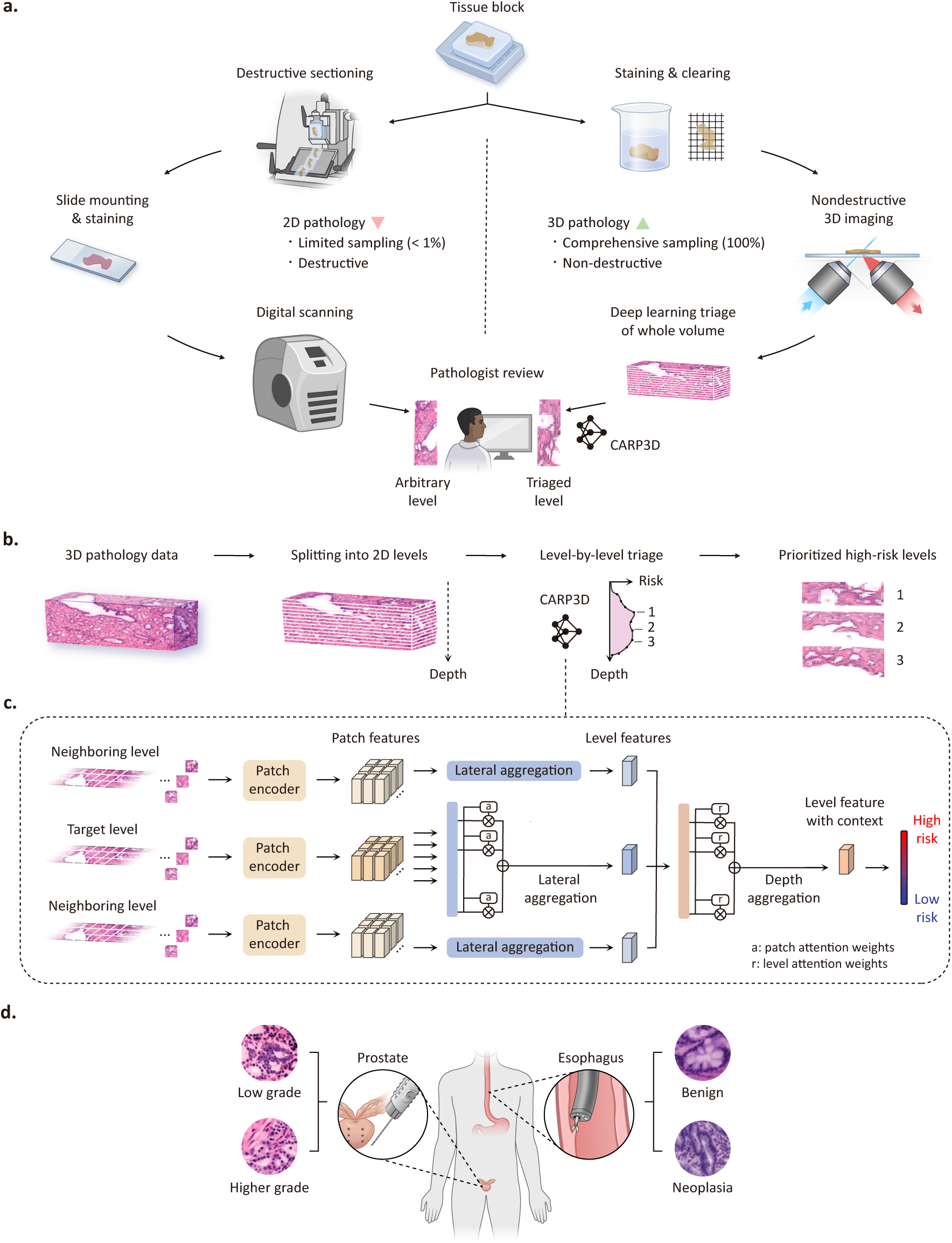
AI-triaged 3D pathology with CARP3D. **(a)** In the current clinical workflow based on 2D histopathology, only a few tissue sections (< 1%) are cut from one arbitrary side of a tissue block for pathologist review. In contrast, CARP3D analyzes the entire tissue in 3D and prioritizes the highest-risk 2D levels for review. **(b)** For triaging with CARP3D, the 3D image dataset is split into a stack of 2D levels. CARP3D then performs a level-by-level analysis to construct a depth-dependent risk profile, from which the highest risk 2D levels may be identified. **(c)** CARP3D processes each 2D level (target) along with a subset of neighboring levels, which are split into 2D patches and featurized using pretrained encoders for 2D histology images. Patch features are first aggregated laterally within each level to construct whole-level feature representations, and then across neighboring levels to incorporate depth context for computing a context-aware 2.5D risk score. **(d)** CARP3D is evaluated for prostate cancer risk stratification and endoscopic screening for esophageal dysplasia/cancer.

Recent AI-driven computational efforts have sought to analyze feature-rich 3D pathology datasets without human intervention. These AI-based methods have either relied on predefined “hand-crafted” features extracted from 3D tissue structures, such as glandular or nuclear morphologies ^30, 31^, or have relied on latent features from deep-learning encoders ^12^ for diagnostic tasks such as patient risk stratification. However, such fully automated approaches will likely require extensive and large-scale validation studies before they can be approved and adopted into clinical practice. On the contrary, a potential low-risk pathway for the early adoption of 3D pathology is to continue to rely on pathologists to render diagnostic determinations. This pathway would utilize an AI framework as a triage tool to identify the highest-risk 2D levels within 3D pathology datasets for time-efficient and effective pathologist review ^2, 32^, in stark contrast to the current clinical practice of sampling tissue sections from arbitrary and sparse locations within a sample (**Fig. 1a**).

An attractive strategy for AI-triaged 3D pathology is to treat 3D datasets as a stack of 2D images and to deploy multiple instance learning (MIL) approaches widely adopted for 2D whole-slide images (H&E slides) ^33–41^. These approaches divide large whole-slide images into smaller image patches, where models are trained to map low-dimensional latent representations of these patches to “weak” slide-level labels that are typically easier to obtain than high-resolution patch-based annotations. However, direct application of 2D MIL approaches to 3D datasets would treat 2D cross sections as independent inputs without explicitly accounting for depth context. This could result in inferior performance for a 3D pathology triage method. Alternatively, a recent 3D MIL approach ^12^ captured volumetric context in 3D image chunks and demonstrated improved performance for patient risk stratification. However, with such 3D MIL approaches, risk predictions are made for entire tissue specimens with no ability to provide fine-grained risk predictions of individual 2D levels within specimens, as is desired to facilitate pathologist review.

In this work, we present context-aware risk prediction in 3D (CARP3D), a 2.5D MIL framework for diagnostic triaging that produces fine-grained risk profiles for all 2D depth levels within a 3D tissue volume, thereby enabling the highest-risk 2D levels within a 3D pathology dataset to be identified for pathologist review. The incorporation of depth context (2.5D MIL) leads to improved diagnostic accuracy with more spatial continuity as a function of depth (i.e. less noise in the risk predictions) in comparison to a 2D MIL baseline. Importantly, our 2.5D method can leverage the powerful foundation models that have recently been developed to encode 2D histopathology images into low-dimensional latent representations ^42–50^. To validate its potential as a general-purpose triage tool for 3D pathology, we implement CARP3D for two clinical scenarios. First, CARP3D performs image triage for prostate cancer risk stratification, ensuring that higher-risk patients receive potentially life-saving treatments while sparing low-risk patients from the side effects of overtreatment. In another application, CARP3D enhances the evaluation of endoscopic biopsies from patients with Barrett’s esophagus for improved detection of dysplasia/cancer, which also has important treatment implications. For preliminary clinical validation, panels of board-certified pathologists reviewed both AI-triaged 3D pathology images and standard 2D histopathology images obtained from the same tissue specimens. These studies indicate that AI-triaged 3D pathology can improve the ability to identify high-risk diseases while optimizing pathologist workloads.

## Results

### Overview of CARP3D pipeline

CARP3D distinguishes itself from previous 3D pathology frameworks ^12, 30, 31^ by adopting a human-in-the-loop approach. Rather than directly predicting clinical outcomes with fully-automated AI, the final diagnosis is rendered by pathologists based on CARP3D-triaged 2D tissue levels that have a familiar H&E-like appearance. Specifically, CARP3D splits a volumetric dataset into a continuous stack of 2D levels and generates a fine-grained risk profile as a function of depth across all levels such that only the highest-risk images are shown to pathologists for time-efficient diagnostic assessments (**Fig. 1b**).

To predict the risk of a 2D level (*target*) within a 3D pathology dataset, CARP3D incorporates morphological information from not only the target level, but also its neighboring levels (**Fig. 1c**). Such a 2.5D approach enables the incorporation of additional contextual information from neighboring depths to enhance level-by-level risk assessments, which also improves spatial continuity (i.e. less noise in the risk assessments) as a function of depth. The target level and its neighboring levels are divided into a set of smaller 2D image patches, which are subsequently compressed into low-dimensional representations through a 2D histopathology encoder pretrained on large numbers of histopathology images ^42–50^. The sets of 2D patch features across the levels are distilled into a single representation via a sequence of two aggregation networks. An attention-based *lateral aggregation* network first averages the 2D image-patch representations of each level into a single whole-level feature vector (latent representation), weighted by the attention score that quantifies each patch’s contribution towards the diagnosis. Subsequently, a *depth aggregation* module integrates contextual information from neighboring levels with the target level, generating a depth-aware *level feature with context* for risk prediction. This aggregation module can emphasize 2D levels that contain diagnostically important information by assigning them with higher weights. This process is repeated for all depth levels across a sample to construct a depth-dependent risk profile for the entire volume.

Incorporating depth context, which is unattainable with conventional 2D histology images, comes with two benefits. First, the added depth context helps to resolve certain ambiguities that arise from viewing complex 3D tissue morphologies as 2D cross sections. In addition, CARP3D learns from the more-diverse tissue morphologies that are seen in neighboring levels, thereby enhancing generalizability and accuracy, especially in a low-data regime.

### Clinical use cases for CARP3D

In the envisioned clinical implementation of CARP3D, the model evaluates all 2D cross sections within 3D pathology datasets and identifies the levels with the highest predicted risk for pathologist review. To this end, we implemented CARP3D for two clinical applications (**Fig. 1d**): prostate cancer risk stratification and esophageal dysplasia/cancer screening.

First, for prostate cancer risk stratification, the low-risk category consists of images exhibiting benign tissue or Grade Group (GG) 1 carcinoma (low-grade prostate cancer) whereas the high-risk category consists of images exhibiting Grade Group 2 or higher carcinoma (higher-grade prostate cancer) ^51, 52^. These two categories of patients typically receive dramatically different treatment recommendations, with low-risk patients often undergoing active surveillance and high-risk patients often receiving curative therapies such as surgery and/or radiation therapy ^53, 54^. The model was developed on 112 cancer-containing core needle biopsies (∼ 10 *mm* × 1 *mm* × 1 *mm*) extracted *ex vivo* from the radical prostatectomy specimens of 54 patients at the University of Washington (UW). The 3D prostate datasets were acquired with a 2nd-generation OTLS microscope ^16^, which achieves an optical resolution of ∼1 *µ*m laterally and ∼4 *µ*m axially. The sampling pitch of the 3D prostate datasets used in prostate model development was ∼1 *µ*m/voxel in all dimensions. A panel of 6 genitourinary (GU) pathologists annotated one or two central levels per biopsy, and the majority opinion among them was used as the ground truth label for the 2D images for model training.

For esophageal dysplasia/cancer screening, the low-risk category consists of images exhibiting normal tissue or non-dysplastic Barrett’s esophagus (NDBE), whereas the high-risk group consists of images exhibiting low-grade dysplasia, high-grade dysplasia, or adenocarcinoma ^55^. Again, accurate identification of higher-risk patients is critical, since they are often recommended to receive curative interventions such as radiofrequency ablation (RFA), endoscopic mucosal resection (EMR) or dissection (EMD), amongst other treatments ^56, 57^. The model was developed on a total of 95 specimens, either endoscopic biopsies or endoscopic mucosal resection (EMR) specimens, obtained from 24 patients at UW. The specimens were imaged using a 3rd-generation OTLS microscope ^14^ with an optical resolution of 0.6 µm laterally and 3 µm axially. The sampling pitch of the 3D esophageal datasets used in this study was 0.4 µm/voxel in all dimensions. Ground-truth labels for selected 2D levels were obtained based on the majority opinion diagnosis of a panel of 3 board-certified pathologists with specialty training in gastrointestinal malignancies.

### Engineering a 2.5D MIL pipeline - CARP3D

To construct an accurate risk-prediction model, which forms the foundation for AI-based triaging, we explored several technical variations of CARP3D for integrating and distilling neighboring depth context (**Fig. 2a**). First, we benchmarked a 2D model using attention-based multiple instance learning (ABMIL) ^34^, a data-efficient approach that aggregates patch features only from one target level. Next, we explored 2.5D MIL strategies that incorporate depth context. CARP3D_N_ with näıve aggregation applies the same ABMIL model to aggregate all patches within the target level and neighboring levels without taking into account spatial location (i.e. all patches are treated equally regardless of lateral or vertical position). In contrast, CARP3D_D→L_ first aggregates patches along the depth dimension using a non-gated attention module and then aggregates laterally using ABMIL. Finally, CARP3D_L→D_ (the default pipeline for CARP3D) first aggregates patch features laterally within each level using ABMIL and then performs depth aggregation of those level features. The depth-aggregation strategy will be described in a later paragraph, including the optimal choice of depth range and the optimal number of levels to aggregate. We employed a leave-one-out cross-validation approach to evaluate model performance. In each fold, images from a single patient were held out for evaluation, while the remaining images were used for model training. The final performance metrics were calculated on predicted risks across the cohort.

**Figure 2:**
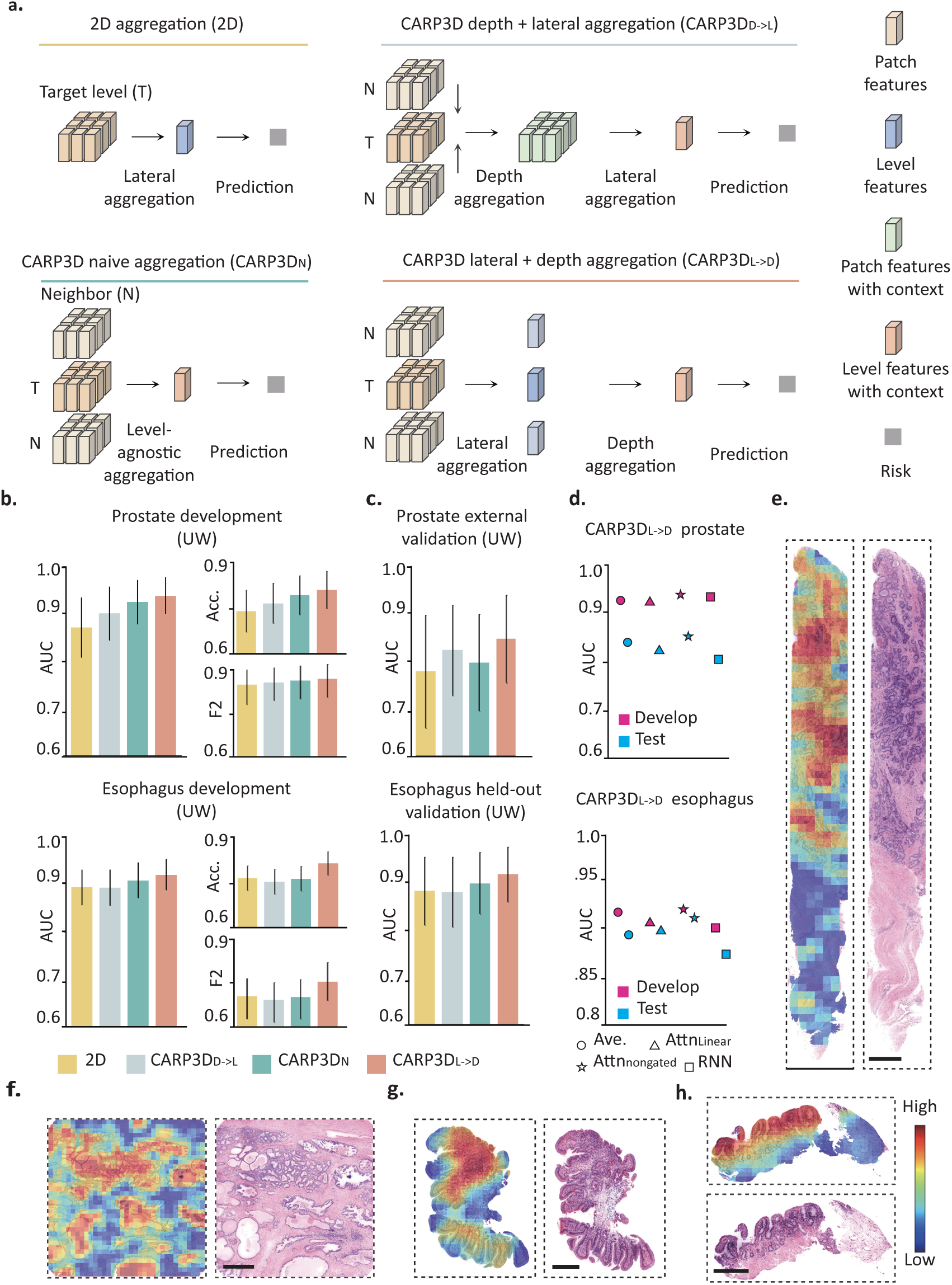
CARP3D analysis of prostate and esophageal specimens. **(a)** Different aggregation strategies: target-level-only aggregation (2D), context-aware depth-and-lateral aggregation (CARP3D_D→L_), context-aware naive aggregation that is agnostic to spatial location (CARP3D_N_), and context-aware lateral-and-depth aggregation (CARP3D_L→D_). **(b)** Cohort-level AUC, balanced accuracy, and F2 scores with 95% confidence intervals for different aggregation strategies applied to prostate and esophageal datasets. **(c)** AUC with 95% confidence intervals for different aggregation strategies evaluated on independent validation cohorts of prostate and esophagus. **(d)** AUC on development cohorts comparing different depth-aggregation strategies: averaging, attention-based aggregation with a linear layer, non-gated attention, and RNN. **(e)-(h)** Attention heatmaps and H&E-analog images from 3D pathology datasets of (**e,f**) prostate and (**g,h**) esophagus. Blue and red colors indicate low and high attention scores, respectively. Scale bars represent 500 *µm*.

The area under the receiver operating characteristic curve (AUC) was calculated to assess the ability of various models to distinguish between low- and high-risk levels (**Fig. 2b**). 95% confidence intervals and statistical differences were calculated based on Delong’s method ^58^. We observe that for prostate cancer risk stratification, all 2.5D approaches outperformed the 2D baseline. Among them, CARP3D_L→D_ achieved the highest AUC of 0.939, exceeding the 2D baseline (AUC 0.871; p < 0.005). The same trend was reflected for esophageal dysplasia/cancer screening, where 2.5D approaches outperformed the 2D baseline and CARP3D_L→D_ demonstrated the largest performance gain (AUC 0.921 vs. 0.895, p < 0.005). The balanced accuracy and F2 score were also calculated for more informative model evaluation during the model-development stage. The F2 scores account for both recall and precision while placing greater emphasis on recall, aligning with a general desire to maximize detection sensitivity for a clinical triage application. The balanced accuracy (Acc.) and F2 scores show the same trend for both clinical use cases, with 2.5D MIL outperforming the 2D baseline and CARP3D_L→D_ achieving the best performance (Acc.: 0.826 vs. 0.750 for prostate and 0.833 vs. 0.779 for esophagus; F20.882 vs. 0.862 for prostate and 0.765 vs. 0.711 for esophagus, **Fig. 2b**). To further validate that the improvement is consistent across different patch encoders, we used a CTransPath model ^42^ pretrained on a large collection of pathology images in a self-supervised manner to conduct the same analyses. The results further confirm that incorporating depth context in 2.5D models enhances performance over the 2D baseline, with CARP3D_L→D_ demonstrating the most significant improvement regardless of the patch encoder used (**Extended Data Figure 1**).

To identify the optimal depth-aggregation strategy for CARP3D_L→D_, we performed ablation studies to compare several architectures: simple averaging (Ave.), attention with a single linear layer (Attn_linear_), non-gated attention (Attn_nongated_) ^34^, and recurrent neural networks (RNN) ^59^. Among these, we observed that non-gated attention was the most effective depth-aggregation strategy during both model development and testing (**Fig. 2d**). Furthermore, the number and spatial range of neighboring depth levels to include were carefully optimized through a matrix of ablation studies for both clinical applications. In short, adding “depth context” is achieved in two ways: incorporating a larger spatial range of information (more morphological diversity) and adding more 2D levels within that range (denser and more highly resolved morphological information). Our findings indicate that performance improves as the depth range increases, but only up to a certain point. Beyond this point, the additional context becomes less helpful since it is less relevant to the target level, leading to a decline in performance (**Extended Data Figure 2c**). We observe that the optimal range for depth aggregation is around 60 *µm* for prostate, which is consistent with the fact that prostate cancer risk stratification (e.g. Gleason grading) is heavily dependent upon relatively large-scale glandular features (architecture) ^53^. On the other hand, the optimal depth-aggregation range for esophageal tissue is around 10 *µm*, which reflects the fact that risk stratification is more dependent on smaller-scale cellular features (cytology) ^60^. These depth ranges are also consistent with previously reported measurements of intra-biopsy variability (spatial heterogeneity) in prostate cancer grades and esophageal metaplasia ^4, 6^. As for the number of levels to include within the optimal range, we found that using a total of three levels, with the target level at the center, yielded the best performance for both clinical use cases. This is potentially due to introducing redundant information when examining too many 2D levels that are closely spaced, which may have a dilutive effect when predicting the risk of the target level. Other key factors, such as false-colored (H&E-like) or dual-channel raw images inputs, choice of patch feature encoders, and lateral aggregation models, were also meticulously optimized to achieve the best performance (**Extended Data Figure 2a, b**).

### Validation of CARP3D with independent test cohorts

To assess the generalization performance of CARP3D, we evaluated the trained model on independent test cohorts for each clinical use case (**Fig. 2c**). For prostate cancer, the test cohort consisted of specimens that were sourced from the University of Pennsylvania (UPenn) and imaged with a newer OTLS microscopy system ^17^ and a slightly modified staining protocol ^13^. Therefore, this test cohort provided a challenging test of the model’s ability to perform well on datasets with non-trivial differences compared with the training dataset. Pathologists labeled three levels per specimen based on 29 specimens from 29 patients. For esophageal dysplasia/cancer, the test cohort of specimens was sourced from the same institution as the training cohort (UW) and was imaged with the same OTLS microscope as the training cohort, but consisted of specimens from patients held out from model development. A total of 77 images from 30 specimens from 5 patients were labeled.

With the independent test cohorts for both clinical applications, 2.5D MIL approaches achieved superior performance to 2D MIL. Notably, CARP3D_L→D_ demonstrated the greatest margin of AUC improvement over the 2D baseline, with an AUC of 0.847 vs 0.779 (p < 0.005) for prostate cancer risk stratification and an AUC of 0.917 vs. 0.882 (p < 0.01) for esophageal dysplasia/cancer screening. The esophageal dysplasia/cancer triage models showed only a slight drop in performance for the independent validation cohort compared with the training cohort. On the other hand, the drop for the prostate model could be attributed to the previously mentioned differences in staining protocols ^13, 30^ and OTLS microscope specifications ^16, 17^ between the training and testing cohorts.

### Histomorphological insights from CARP3D

Encouraged by the strong performance of CARP3D, we sought to derive interpretable insights from the predictions. Specifically, each 2D patch receives an attention score from the lateral aggregation module based on its relative contribution to the final prediction. We can visualize these scores in the form of heatmaps overlaid on the tissue images. As illustrated in **Fig. 2e-h, Extended Data Fig. 3, 4, Supplementary Video 1, 2**, in the case of prostate cancer, higher attention is predominantly focused on glandular structures, aligning with the fact that pathologists rely on glandular architecture for prognostic Gleason grading. In the higher-grade prostate cancer image (**Fig. 2e**), attention is concentrated on regions with fused glands, characteristic of Gleason pattern 4. In contrast, in the low-grade prostate cancer image (**Fig. 2f**), the model highlights distinct well-formed cancer glands (Gleason pattern 3) and large benign glands. For esophageal images (**Fig. 2g**), the model places high attention on crowded and fused glands, a hallmark of high-grade dysplasia. In comparison, for low-grade dysplasia, attention is primarily focused on the epithelial surface, where nuclei appear dark, elongated, and stratified (**Fig. 2h**). During depth aggregation, certain levels consistently receive higher attention scores, whether acting as the target level or as a “neighboring level,” suggesting that these levels hold important features for rendering final predictions.

### CARP3D reveals tissue heterogeneity with depth

Following the optimization of critical design parameters, we deployed CARP3D to generate predicted risk profiles of volumetric samples, resulting in more stable and accurate profiles compared to the 2D MIL approach (**Extended Data Fig. 5**). This provides insights into the degree of heterogeneity in tissue morphologies and their associated risks as a function of depth. More specifically, we generated risk predictions, ranging from 0 to 1 for lowest to highest risk, for 2D levels spaced at 10-*µm* depth intervals within 3D datasets from independent test cohorts (**Figs. 3a-d**). Three representative 2D levels (top, middle, and bottom) are shown from each example 3D dataset. For each of these levels, the 2D image patches that receive the highest attention scores are displayed. Finally, for the latent features from the 15 top-attended image patches per level, the first two principal components (PC) are plotted, color-coded based on their depth level.

**Figure 3:**
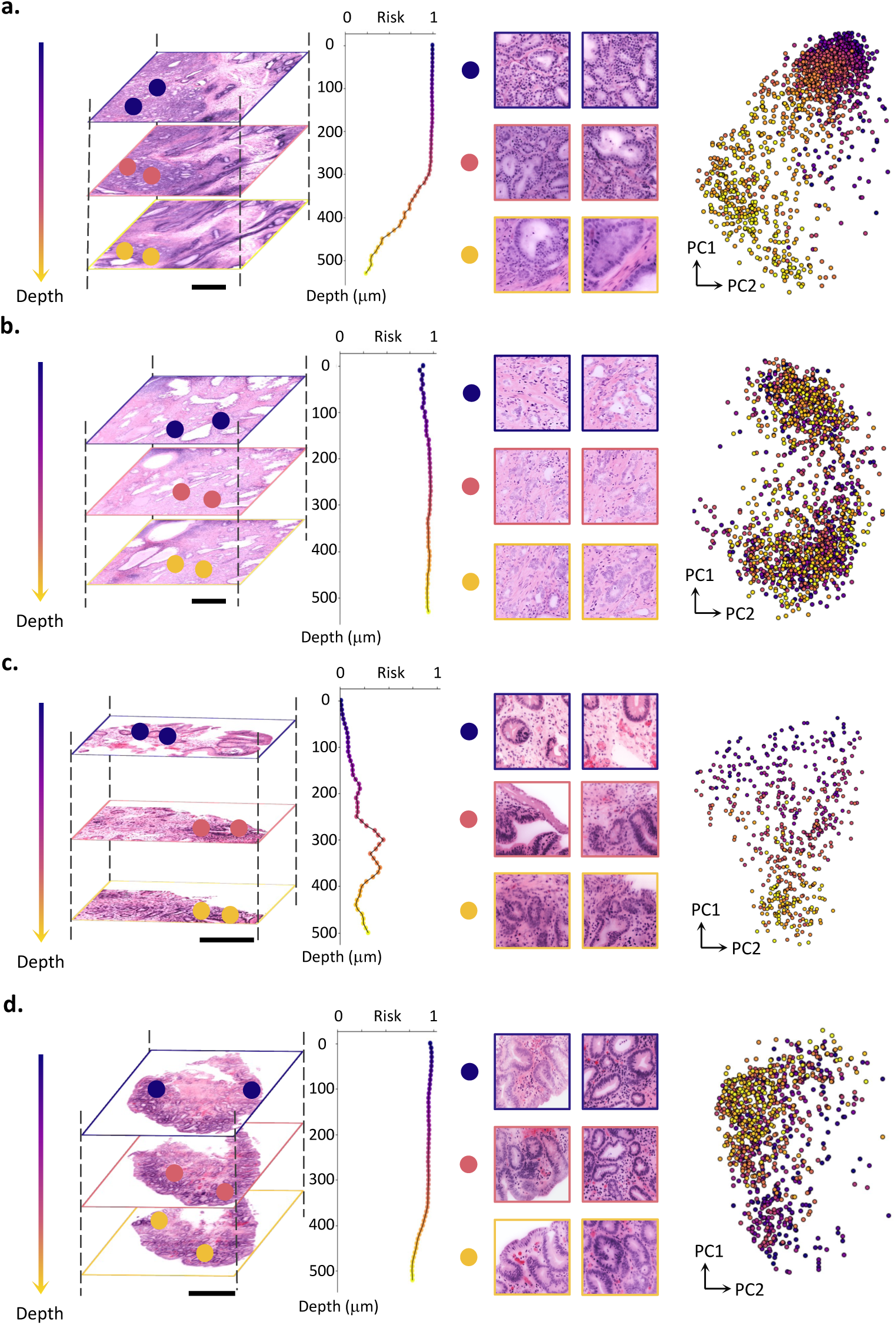
Depth-dependent risk predictions for 3D pathology datasets with CARP3D. Visualization of three representative 2D levels (top, middle, bottom) from a 3D tissue volume, along with the corresponding risk profile as a function of depth. The two top-attended patches are displayed for each 2D level. The first two principal components (PCs) are shown for the featurized representations of the 15 top-attended patches across all levels, color-coded by depth to reveal depth-associated clusters. Datasets with varying risk scores, corresponding to heterogeneous morphologies along the depth axis, are displayed for **(a)** prostate and **(c)** esophagus. Datasets with relatively uniform risk scores, corresponding to homogeneous morphologies along the depth axis, are displayed for **(b)** prostate and **(d)** esophagus. Scale bars represent 500 *µm*.

**Fig. 3a** and **3b** illustrate variations in risk as a function of depth for prostate datasets. In **Fig. 3a**, the top portion of the tissue volume is predicted to have a high probability of containing higher-grade prostate cancer, with decreasing risk towards the bottom portion of the volume. The top and middle 2D levels, which contain top-attended patches, reveal areas of fused and crowded glands, a hallmark of higher-grade prostate cancer (i.e. Gleason pattern 4). In contrast, the bottom level contains larger benign glands and fully formed Gleason pattern 3 glands (low-grade prostate cancer). The corresponding PC plot shows that the top-attended patches from different levels form distinct clusters, further illustrating the high heterogeneity (in latent features) across depth in this prostate specimen. In **Fig. 3b**, the 3D dataset exhibits high-risk predictions across all levels. Example 2D levels and patches with top attention scores show poorly formed glands (Gleason pattern 4), a morphology associated with higher-grade prostate cancer. Unlike **Fig. 3a**, the PC plot indicates that top-attended patches from different levels have latent features that are not distinctly clustered, suggesting that the tissue morphologies are more homogeneous as a function of depth.

**Fig. 3c** and **3d** show risk variations for 3D esophageal datasets. In **Fig. 3c**, the risk scores fluctuate with depth, with the top level and its highly attended patches showing goblet cells with peripheral nuclei, characteristics of non-dysplastic Barrett’s esophagus, and the middle and bottom levels exhibiting elongated and stratified nuclei extending to the surface, a key feature of low-grade dysplasia. **Fig. 3d** shows high-risk morphologies throughout the volume, with example 2D levels and their representative patches revealing low-grade dysplasia. The corresponding PC plot for **Fig. 3c** shows separate clusters of top-attended patches in feature space, reflecting higher morphological heterogeneity. On the contrary, the PC plot for **Fig. 3d** shows a mixed cluster of features, suggesting more homogeneous morphologies as a function of depth.

### Comparison with standard-of-care 2D histopathology

We conducted a study to evaluate whether an AI-triaged workflow improves the ability of pathologists to detect high-risk diseases compared to standard-of-care 2D histopathology. In this preliminary clinical validation study, independent cohorts of specimens (59 prostate samples and 30 esophagus samples) were first imaged non-destructively in 3D before sectioning them as standard H&E histology slides. Pathologists (five for prostate and three for esophagus) reviewed the highest-risk levels identified within 3D pathology datasets by CARP3D, as well as conventional 2D slides prepared from the same biopsy specimens, with a one-month washout period between these assessments (**Fig. 4a**). Following the standard practice for assessing prostate biopsies at most institutions, three levels per biopsy were presented to pathologists for both the 3D-triaged and 2D workflows. The 3D-triaged levels were from AI-selected locations within the whole biopsy in order of risk, while ensuring a minimum separation between triaged levels, whereas the slide-based H&E histology levels were sectioned from one arbitrary side of the biopsy and spaced by 20 microns per standard practice. For esophageal dysplasia/cancer screening, 16 consecutive sections were obtained per biopsy for 2D slide-based evaluation, following the standard practice at UW. However, for AI-triaged 3D pathology, CARP3D was used to select the eight highest-risk levels with a minimum distance maintained between triaged levels.

**Figure 4:**
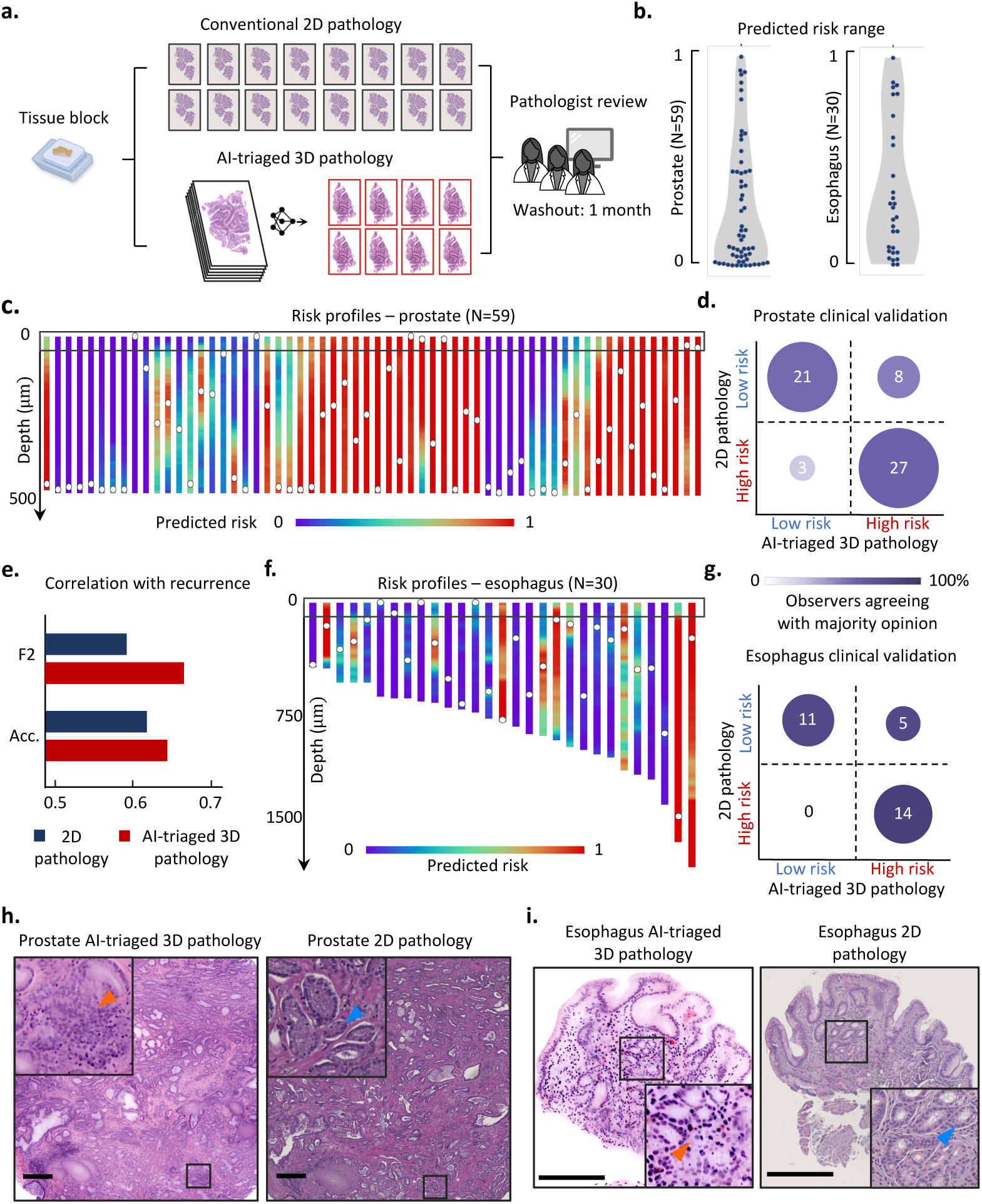
Clinical validation of AI-triaged 3D pathology. **(a)** Panels of pathologists reviewed biopsy images that were generated with both AI-triaged 3D pathology (CARP3D) and conventional 2D histopathology. **(b)** Range of predicted risk scores for each sample in the prostate (*N* = 59) and esophagus (*N* = 30) validation cohorts. **(c)** Depth-dependent risk profiles for all prostate samples. The white dot indicates the highest-risk depth level for each sample. The box indicates the simulated depth range (40*µm*) typically evaluated with 2D histopathology. **(d)** Pathologist diagnoses of prostate cancer biopsies based on 2D histopathology vs. AI-triaged 3D pathology (CARP3D). The assignment of categories for each specimen is based on the majority-opinion diagnostic label for each modality. The circle size and color represent the number of cases and the percentage of observers agreeing with the categorizations shown, respectively. **(e)** Correlation between pathologist-diagnosed risk category (CARP3D or 2D histopathology) and 5-year biochemical recurrence outcomes for the prostate cohort. **(f)** Same as (c) for esophageal samples. The box indicates the simulated depth range of 80 *µm* (16 consecutive tissue sections) typically evaluated at UW. **(g)** Same as (d) for esophageal dysplasia/cancer screening **(h)** Prostate example upgraded with AI-triaged 3D pathology, where fused glands (orange) are identified in 3D and only discrete glands (blue) are seen in 2D. **(i)** Esophagus example upgraded with AI-triaged 3D pathology, where cribriform glands with prominent mitoses are observed in 3D while only non-dysplastic Barrett’s esophagus is seen in 2D. Scale bars represent 300 *µm*.

We quantified the predicted risk range across depth, which was defined as the difference between the 1 percentile and the 99 percentile of risk scores across all 2D levels of each sample. Examining all samples in the validation cohorts, shown in **Fig. 4b**, reveals that 9 of 30 esophageal samples and 11 of 59 prostate samples exhibit substantial risk variance (> 0.5) across depth. To further investigate this variation, we visualized the risk profiles as a function of depth for the prostate (**Fig. 4c**) and esophagus cohorts (**Fig. 4f**), where red and blue indicate high and low risk, respectively. We observe that the depth of the level with the highest predicted risk for each sample (white dot) varies greatly across the cohorts, with only seven samples (11.9% of the prostate cohort) and four samples (13.3% of the esophageal cohort) containing the highest-risk level within the simulated depth range that is typically evaluated with standard 2D histopathology (black box, 40 *µm* for prostate biopsies and 80 *µm* for esophageal specimens). This suggests that AI-triaged 3D pathology, which can identify the highest-risk levels regardless of depth, can reduce misdiagnoses arising from sampling sparse 2D sections from an arbitrary side of a large tissue specimen.

Upon aggregating reader-study results, we computed the Fleiss’ kappa metric ^61^ to assess inter-observer diagnostic agreement for all samples in the cohorts. Pathologist agreements were comparable between AI-triaged 3D pathology (0.357 for prostate and 0.770 for esophagus) and conventional 2D pathology (0.415 for prostate and 0.764 for esophagus). This suggests that the visual interpretability between false-colored H&E-like 3D pathology images and real 2D H&E histology images is comparable, as has been reported in prior studies ^3, 12^.

**Fig. 4d** compares low-versus high-risk cases identified through AI-triaged 3D pathology and conventional 2D pathology. This is based on the diagnostic label provided by the majority of pathologists for each modality. The circle size reflects the number of cases in each category and the darkness of the color indicates the percentage of pathologists who agree with the categorization of the cases as displayed in **Fig. 4d**. For prostate cancer, pathologists upgraded eight cases from low-grade to higher-grade cancer using AI-triaged 3D pathology (**Fig. 4h, Extended Data Fig. 6**). Three cases were also downgraded based on 3D pathology. However, these three cases exhibited low agreement with the downgrading categorization, suggesting diagnostic ambiguity as a contributing factor (**Extended Data Fig. 7**).

The improved classification of high- and low-risk cases with AI triage is also indicated by an improved accuracy for predicting five-year biochemical recurrence (BCR) outcomes. Here, we used the pathologist diagnoses (low- vs. high-risk) to predict a binary outcome of BCR-free survival within five years of surgery ^51, 62^. We observe that AI-triaged 3D pathology yields higher balanced accuracy (0.633 vs. 0.614) and F2 scores (0.654 vs. 0.588), supporting its potential for improved risk stratification (**Fig. 4e**). The distribution of diagnostic labels provided by the panel of pathologists, and the majority opinion for each prostate sample, is shown in **Extended Data Table 1**.

For esophageal dysplasia/cancer screening, pathologists upgraded five cases after reviewing AI-triaged 3D pathology, with no cases being downgraded (**Fig. 4g**). These upgraded cases all have strong agreement between individual pathologists and with the majority opinion. Notably, our analyses reveal that pathologists are able to identify more dysplastic/cancer samples while reviewing only half the number of images (8 vs.16). To examine how far we could reduce pathologist workloads with AI triaging, we asked pathologists to evaluate only three levels per sample after a one-month washout period from when they evaluated eight levels (**Extended Data Fig. 8a**). In this case, fewer neoplastic cases (three levels: 14 vs. eight levels: 17) were identified. This suggests that the eight levels identified by CARP3D strikes a balance between optimal detection and reduced clinical workload for the diagnosis of esophageal dysplasia/cancer via AI-triaged 3D pathology. The distribution of diagnostic labels provided by the panel of pathologists, and the majority opinion for each esophagus sample, is shown in **Extended Data Table 2**.

Examples of upgraded cases demonstrate that the triaged levels contain morphologies that are not captured with standard 2D histopathology (**Fig. 4h, i, Extended Data Fig. 6**). For prostate, fused glands (a variant of Gleason Pattern 4) are observed in AI-triaged 3D pathology images while 2D slides from the same sample only display distinct well-formed cancer glands (Gleason Pattern 3). For esophagus, an AI-triaged 3D pathology image reveals cribriform glandular architecture with prominent mitosis, indicating high-grade dysplasia. In contrast, a 2D slide from the same specimen displays distinct well-formed glands with goblet cells, consistent with non-dysplastic Barrett’s esophagus (**Fig. 4i, Extended Data Fig. 8b, c**).

## Discussion

There is a growing interest in non-destructive 3D pathology, in which valuable clinical specimens (e.g. biopsies) can be comprehensively imaged at high resolution while maintaining those specimens entirely for downstream assays. This is in contrast to conventional histology, which requires destructive sectioning of specimens onto glass slides. With the potential for fully automated tissue preparation (no sectioning) and slide-free digital image acquisition, 3D pathology could also provide efficiency advantages for clinical labs in comparison to 2D workflows that require manual slide preparations by experienced histotechnicians ^1^.

Fully computational AI-based analyses of 3D pathology datasets are being explored, with early studies showing clear advantages in comparison to 2D computational analyses ^2, 12, 30, 31^. However, for early and rapid clinical adoption and regulatory approval, a potential low-risk strategy would be to continue to rely upon pathologists to render final diagnostic determinations. Due to the immense size and information content within 3D pathology datasets, a method for AI triage is needed, in which the highest-risk 2D levels (images) within a 3D pathology dataset can be presented to pathologists for accurate and time-efficient review.

We have developed CARP3D, a 2.5D multiple instance learning (MIL) framework that incorporates morphological context from an optimal subset of neighboring depth levels to improve risk predictions for all 2D levels within a 3D pathology dataset. This provides a depth-dependent risk profile for clinical specimens, from which the highest-risk levels may be identified for pathologist review. Based on optimized design elements and parameters within the CARP3D pipeline, clinical validation studies are presented for two use cases: prostate cancer risk stratification and esophageal dysplasia/cancer screening. These studies show that CARP3D-triaged 3D pathology enables pathologists to better detect high-risk disease than conventional 2D pathology, with the added benefit of optimizing pathologist workloads.

Compared to “gold-standard” biopsy analysis of sparse tissue sections extracted from an arbitrary side of each biopsy, the ability to identify the optimal levels for pathologists to review across whole specimens could be transformative for patient care. Our study reveals that biopsies can contain a high degree of spatial heterogeneity in terms of histomorphology, and that the highest-risk regions can be missed by standard 2D histology due to the limited depth range that is evaluated.

AI-triaged 3D pathology represents a general-purpose pipeline that is applicable to a broad range of diagnostic tasks in addition to the clinical applications explored here. Nondestructive 3D imaging platforms — such as OTLS microscopy ^14–18^, holotomography ^63–65^, and micro-CT ^19, 66, 67^ — can capture high-resolution images of specimens from diverse organs of various shapes and sizes. These platforms generate large volumes of data that support both AI-driven analysis and human interpretation. While this study focused on identifying the best depth levels within relatively small biopsies for pathologists to view as 2D images, AI triage of larger specimens, such as from surgically excised tissues ^68–70^, could facilitate the identification of 2D regions of interest that are localized both laterally and in depth for efficient and effective pathologist review. Emerging AI foundation models ^45–50, 71^ will accelerate the rapid development of task-specific triage models for diverse applications of 3D pathology.

Due to imperfect tissue clearing with the simple and rapid protocols used in this study, image quality deteriorates with increasing imaging depth ^13^. In this study, relatively high-quality imaging was possible at a depth range of 0.5 to 2 millimeters, which corresponds to the diameter/size of most biopsies. More sophisticated tissue-clearing protocols ^72, 73^, advanced imaging systems with aberration and scattering mitigation ^74–76^, and robust denoising algorithms ^77, 78^ may facilitate the imaging of thicker or more highly scattering/aberrating specimens, if needed. There are subtle differences between 3D pathology images and conventional histology, such as the lack of retraction or dehydration artifacts and cracks from physical sectioning. Large-scale studies are needed to more rigorously assess and calibrate the human interpretation of 3D pathology datasets and to further refine the diagnostic determinations and treatment recommendations made with AI-triaged 3D pathology images. This is especially important since current clinical management decisions have been developed and optimized based on traditional slide-based 2D histopathology, as imperfect as it may be.

Lastly, while the clinical cohorts presented in this study are among the most substantial to date in the emerging field of 3D pathology, datasets comprising more diverse clinical specimens and tissue-preparation protocols are needed to further enhance the robustness and generalizability of AI models, including more optimal patch feature encoders and MIL models. Furthermore, ongoing efforts aim to develop triage models capable of finer-grained risk stratification (e.g. multi-class grading/staging) to facilitate more nuanced clinical decision support. Integration with multimodal data sources, such as molecular profiles ^79–81^, immunohistochemical staining ^82^, and clinical reports ^44, 83^, can also enable more informative, explainable, and accurate analyses.

In summary, CARP3D enhances the identification of high-risk 2D levels in 3D pathology datasets by efficiently and effectively leveraging depth context. Based on CARP3D, initial clinical validation studies suggest that diagnostic tasks can be improved with AI-triaged 3D pathology even while reducing pathologist workloads.

## Online Methods

### Dataset description

#### Prostate development cohort

Archived formalin-fixed, paraffin-embedded (FFPE) specimens were collected from the prostatectomies of 54 prostate cancer patients treated at the University of Washington Medical Center (UWMC). For each case, FFPE blocks were identified corresponding to the six prostate regions targeted by urologists when performing standard sextant (6-core) and 12-core biopsy procedures. From each of the six blocks, one simulated core-needle biopsy (roughly 1 mm × 1 mm × 15 mm) was extracted. Following OTLS imaging ^16^, 112 biopsies were identified as containing cancer by a board-certified pathologist. For the 3D pathology dataset of each cancer-containing biopsy, one or two representative 2D levels (typically at the center depth) were selected for annotations. A panel of six pathologists reviewed the images and a majority opinion on low-grade (Grade Group 1) versus higher-grade (Grade Group > 1) prostate cancer was established at 121 2D levels for model development (53 low-grade, 68 higher-grade).

#### Prostate validation cohort

Archived FFPE proctectomy specimens from 29 patients were obtained from the University of Pennsylvania (UPenn). A board-certified pathologist reviewed the corresponding slides to identify cancerous regions, from which one 3-mm diameter punch biopsy (> 0.5-mm thick) was extracted per case. A panel of three pathologists reviewed three 2D levels, spaced by 20 *µm*, near the center of each 3D dataset to form the test set for benchmarking model generalizability (63 low-grade, 18 higher-grade). For clinical validation, the cohort was extended to 59 punch biopsies from 59 patients, and a panel of five GU pathologists performed diagnostic evaluations.

#### Esophagus development cohort

Archived esophageal biopsy and endoscopic mucosal resection (EMR) specimens from 24 patients were collected at UWMC. A total of 95 specimens were imaged in 3D and multiple 2D levels were extracted from each 3D dataset to better capture dysplasia, which is often localized. A panel of three pathologists labeled 334 2D levels for model development, where 64 contained dysplasia/cancer and 270 levels were benign.

#### Esophagus validation cohort

An independent validation cohort was assembled from 30 specimens collected from five patients at UWMC. A total of 77 levels were extracted from 3D datasets of these specimens (typically near the central depth), which were labeled by a panel of three pathologists (34 benign, 43 dysplasia/cancer). For clinical validation, all 30 specimens were included and a panel of 3 pathologists performed diagnostic evaluations.

### Tissue preparation

#### Deparaffinization

FFPE blocks were first deparaffinized in an oven at 75 °C for 1 hour to melt the outer paraffin and then placed in xylene at 65°C for 48 hours. Subsequently, the specimens were washed in 100% ethanol (EtOH) twice, for 1 hour each, to remove residual xylene.

#### Staining

For the prostate development cohort, core-needle biopsies cut from each FFPE block were rehydrated partially in 70% EtOH for 1 hour. Each biopsy was then stained for 48 hours in 70% ethanol (pH 4) using a 1:200 dilution of Eosin-Y and a 1:500 dilution of TO-PRO-3 Iodide. Staining was performed at room temperature (RT) in 0.5 mL tubes with gentle agitation.

The prostate validation cohort was stained using a recently updated protocol for improved quality ^13^. After deparaffinization, specimens were washed twice in 100% EtOH (1 hour per round), followed by rehydration in 70% EtOH for 3 hours. Each specimen was then incubated in 1.5 mL of staining solution (1:500 TO-PRO-3 and 1:100 Eosin-Y) for 48 hours with gentle shaking at RT.

The esophagus development and validation cohorts were stained using a customized protocol optimized for esophageal tissue ^3^. Specimens were first washed twice in 100% EtOH for 1 hour each, then incubated sequentially in 75%, 50%, and 25% EtOH (1 hour per step). They were subsequently washed in phosphate-buffered saline (PBS) for 1 hour, followed by incubation in 1:500 phosphate-buffered saline Triton (PBST). Staining was performed overnight at RT using a 1:1500 dilution of TO-PRO-3 in PBST. Afterward, tissues were washed in PBST and PBS (1 hour each), then gradually dehydrated in 25%, 50%, and 70% EtOH. Finally, specimens were incubated overnight in 70% EtOH (pH 4, adjusted with hydrochloric acid) with a 1:2000 dilution of Eosin-Y at RT.

#### Clearing

After staining, specimens were dehydrated twice in 100% ethanol for 2 hours each. The biopsies were then optically cleared (refractive index = 1.56) by incubation in ethyl cinnamate (ECi) for 8 hours at RT before imaging.

### 3D imaging with OTLS microscopy

All specimens were imaged using custom-built OTLS systems^14, 16, 17^. Tissues were placed on flat, index-matched sample holders (Hivex, 200-um thick) and imaged from below through ECi immersion media (n=1.56). A thin light sheet illuminated the specimens at a 45-degree angle. Fluorescent signals were collected by the objective lens and relayed to a high-speed sCMOS camera. The scanning stage moved the sample along the x, y, and z axes, during which adjacent images were acquired and saved to form a continuous 3D dataset.

The prostate development cohort was previously imaged using a 2nd-generation OTLS system ^16^, which provides submicron lateral resolution and 3.5 *µm* axial resolution. Imaging was conducted at near-Nyquist sampling (0.44 *µm*/voxel). Nuclear fluorophores were excited using a 638 *Nm* wavelength and eosin was excited at 488 *Nm*.

The prostate validation cohort, sourced from UPenn, was imaged using a 4th-generation OTLS system ^17^ with improved aberration correction and optical resolution (< 1 *µm* lateral, ∼ 2.9 *µm* axial). Imaging was performed at a sampling pitch of 0.37*µm*/voxel. Nuclear fluorophores were excited at 638 *Nm* while eosin was excited at 561 *Nm* to minimize tissue scattering and thereby improve imaging depth.

To better resolve the sub-cellular details that are important for esophageal tissue diagnostics, the esophagus development and validation cohorts were imaged using a high-resolution 3rd-generation OTLS system ^14^. This system achieved ∼ 0.6 *µm* lateral and ∼ 2.75 *µm* axial resolution, with a voxel sampling pitch of 0.21 *µm*. Nuclear fluorophores were excited at 638 *Nm* and eosin was excited at 488 *Nm*.

### 3D data preprocessing

The 3D datasets, acquired in the format of tiles of 2D images, are first stitched and then fused at 2x downsampling by the Bigstitcher software ^26^ (for all prostate samples) or a self-developed rapid stitching software ^84^ (for all esophagus samples). The downsampling enables efficient data processing in subsequent steps, while maintaining sufficient resolution for prostate cancer grading (which is based on glandular morphology) and esophageal dysplasia/cancer screening (with near-Nyquist sampling). For the prostate development cohort, each volumetric dataset (∼ 1*mm* × 1*mm* × 15*mm*) amounts to ∼ 1000 × 1000 × 15000 voxels. For the prostate validation cohort, the central square of each cylindrical punch (∼ 2*mm* wide and on average ∼ 0.5*mm* in depth) is cropped, which equates to ∼ 3500 × 3500 × 800 voxels. For the esophagus development and validation cohorts, endoscopic biopsies or EMR specimens vary in size from around 1*mm* to 5 *mm* (2500 – 12000 voxels) in lateral extent and from ∼ 0.5 to ∼ 2 *mm* (1000 to 5000 voxels) in depth.

The two-channel fluorescence images were false-colored to mimic the appearance of H&E images using an open-source package based on the Beer-Lambert law of absorption^27^. We represent each false-colored 3D pathology data as a stack of 2D images 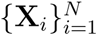, where **X***_i_* ∈ ℝ*^W^* ^×^*^H^* with *W* and *H* denoting width and height, and *N* referring to the number of 2D depth levels in 3D data.

### Model architecture

For a target level in a 3D pathology dataset **X***_k_* where *k* ∈ {1*, …, N* }, we aim to predict its label for the risk category *y_k_*. Additional depth context from its neighboring levels spaced by an interval of *d*, denoted as 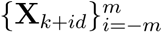, is leveraged to enhance the predictions. Tissue regions within each of the levels are first segmented by Otsu threholding, and are then split into small patches. We denote the set of patches forming an image **X***_i_* as 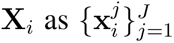 where 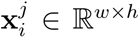, with *w* and *h* representing lateral dimensions of the patches and *J* indicating the total number of patches in **X***_i_*.

#### Patch feature encoding

A pretrained feature encoder *f*_enc_ extracts a low-dimensional and representative feature 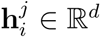 from each patch 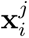, such that 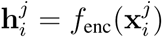.

Here, CONCH ^71^ is selected as the patch feature encoder for several reasons. As a foundation model pretrained on millions of histology–text pairs, CONCH provides rich and robust feature representations of histomorphologies for downstream prediction tasks. Additionally, its training corpus includes non-histology images such as special stains, which may improve the ability to generalize to our OTLS datasets. Finally, CONCH produces lower-dimensional embeddings (*d* = 512) compared to other histopathology foundation models that typically produce embeddings of *d* = 768 or higher, thus enabling CONCH to produce simpler models that are better suited for low-data regimes (i.e. to avoid overfitting).

To address the domain discrepancy arising from applying encoders, pretrained on standard 2D H&E histology images, to OTLS-generated images, we apply a fully-connected layer with ReLU nonlinearity to 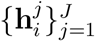 to adapt original patch features to better align with the target domain 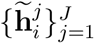. Specifically, the more domain-specific features, 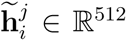, are computed based on 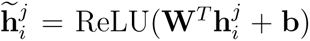, where **W** ∈ ℝ*^d^*^×512^ and **b** ∈ ℝ^512^.

#### Lateral aggregation

The fine-tuned patch features 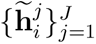 are aggregated by an attention network into a whole-level feature *z_i_* ∈ ℝ^512^, where the attention score 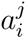 computed for corresponding patch feature 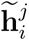 reflects its importance to the final prediction. The attention network is parameterized by three matrices **V** ∈ ℝ^512×256^, **U** ∈ ℝ^512×256^, and **W** ∈ ℝ^256×1^. The calculation follows a gated attention mechanism, which enables the capture of complex relationships among patches.

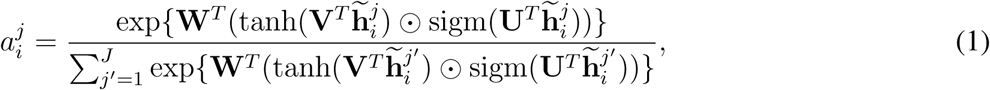

where 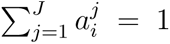, tanh and sigm denote hyperbolic tangent and sigmoid function respectively, and ⊙ denotes element-wise multiplication operation. The individual patch features are weight-averaged by their corresponding attention scores, resulting in an aggregated feature **z***_i_*,

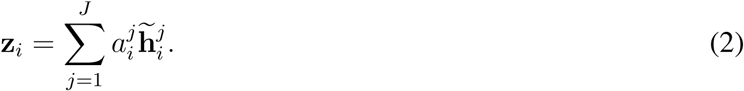

#### Depth aggregation

While **z***_i_* alone could yield a prediction, we show that leveraging contextual information from neighboring levels can improve the prediction of the target level **X***_k_*. A total of *m* levels above and below **X***_k_* with a spacing of *d* between levels, i.e., 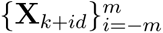 are condensed to a set of level features 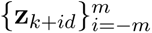.

Similar to lateral aggregation, we apply an attention module to aggregate a target level and its neighboring levels 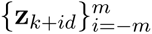. However, since this module operates on a small number of level features that are more discriminative than patch features, a simpler non-gated attention mechanism is used to calculate depth attention scores on each level, denoted as 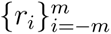. Specifically, we introduce **K** ∈ ℝ^512×256^ and **L** ∈ ℝ^256×1^ to aggregate level features across depth and to construct a context-aware feature -**z̃***_k_*.

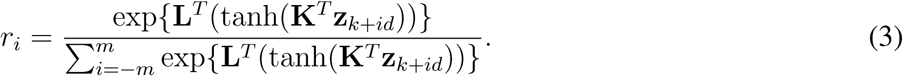

The target-level feature with context **z̃***_k_* is then calculated as the weighted average of a set of neighboring level features,

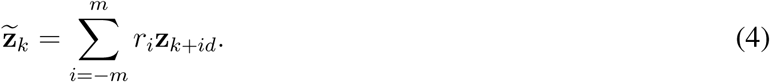

Here *r_i_*refers to the weight of **z***_k_*_+*id*_ and reflects its relevance to the final prediction, summed up to 1.

#### Classification module

As a final step, the prediction *ŷ_k_* on **X***_k_* is calculated as follows. Here **C** is the classification layer, **C** ∈ ℝ^512×^*^N^*, and *N* is the number of risk classes. *N* = 2 for this study.

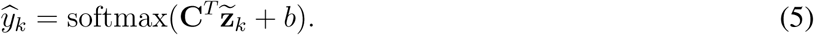

### Ablation on aggregation strategies

#### 2D aggregation

As the baseline, only patch features within the target level are aggregated by the above gated attention mechanism described in **Eqs. 1** and **2**, producing **z***_i_* for classification in **Eq. 5**.

#### Naive aggregation (CARP3D*_N_*)

To aggregate depth context, naive aggregation employs the above gated attention mechanism to aggregate patch features across the target level and neighboring levels in one step, agnostic to their depth positions. The attention scores are calculated as follows.

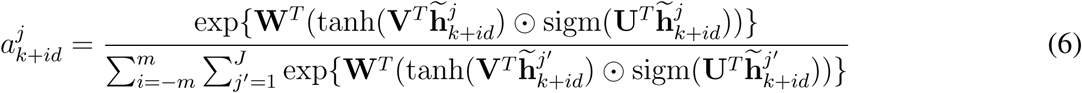

#### Depth and lateral aggregation (CARP3D*_D_*_→*L*_)

Another strategy is to first aggregate patch features along the depth direction to prioritize the learning of vertical dependencies, producing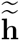, such that

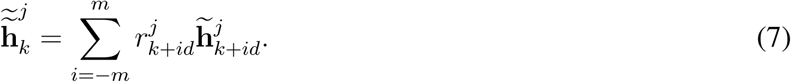

where 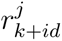 corresponds to the depth attention weighted on patch features of neighboring levels 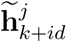. These weights are computed using a non-gated attention mechanism.

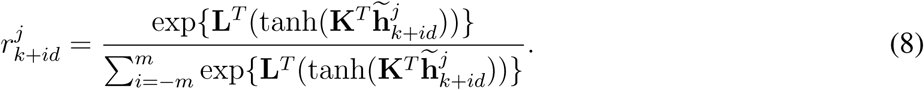

Subsequently, patch features (containing depth context) within the target level are aggregated laterally based on the above non-gated attention mechanism as **Eq. 1** to construct a context-aware feature for final risk predictions in **Eq. 5**

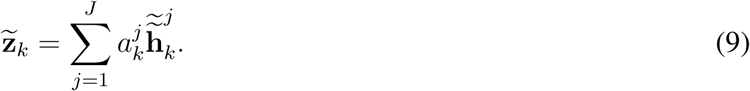

where 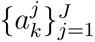 are calculated as follows

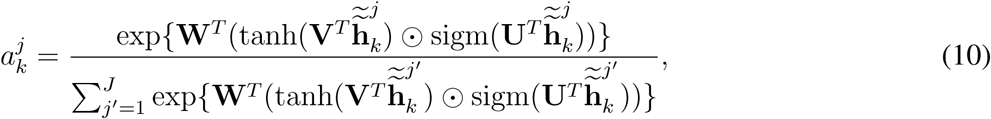

### Depth aggregation modules for CARP3D*_L_*_→_*_D_*

#### Averaging

We further explored different modules to aggregate neighboring level features (i.e. “depth aggregation”), with the most intuitive strategy being averaging across depth.

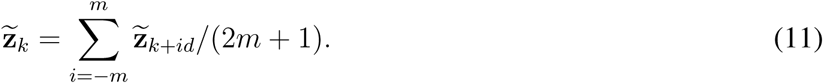

#### Linear attention

Similar to **Eq. 7**, the target level feature with context **z̃***_k_* is computed by a weighted averaging of neighboring level features, allowing more important 2D levels to be dynamically weighted. However, instead of non-gated attention, we used a simple linear layer to produce the set of weights 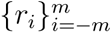. Specifically, we introduce **P** ∈ ℝ^512^ such that

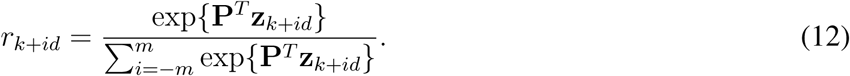

#### Recurrent unit

Finally, we experimented with using recurrent units to aggregate depth context, as they have been shown to have value for capturing sequential information. Recurrent units aggregate image sequences 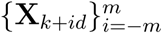 in a bi-directional manner, producing **hid** as hidden state features with context information separately from above and below the target level,

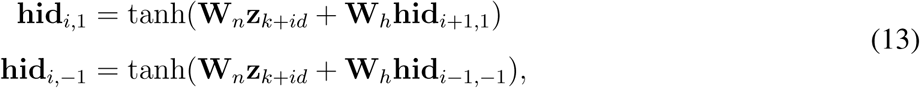

where **W***_N_,* **W***_h_* ∈ ℝ^512×512^ are weight matrices and +1 or −1 denotes the direction of integration. We concatenate the hidden state features at level *k* to form -**z***_k_*, where -**z***_k_* = **hid***_k,_*_1_ ⊕ **hid***_k,_*_−1_.

### Training and inference

#### Training

During the training phase, we employed a leave-one-out cross-validation strategy to benchmark model performance. In each fold, 2D levels from one patient were held out as the testing set, while the remaining images were used to train the model. The metrics were calculated at a cohort level across predictions on all folds.

For the prostate model, images were split into 256 × 256 patches (∼ 200*µm* per side) and were subsequently encoded into 512-dimensional feature vectors through CONCH. Both 2D and 2.5D models were trained using the Adam optimizer with a constant learning rate of 2 × 10^−4^ and a weight decay of 10^−5^ to iteratively minimize the cross-entropy loss. A weighted sampling strategy was applied during training to balance the class distribution within each epoch. Models were trained for 50, 100, and 150 epochs to select the best-performing model based on AUC metrics.

For 2.5D analysis, we explored incorporating neighboring levels located at distances (*md*) of 20, 40, 60, and 80 *µm* above and below the target level, with the total number of levels selected from *m* ∈ {1, 2, 4}. A single pair of neighboring levels (*m* = 1) spaced 60 *µm* apart yielded optimal performance.

For the esophagus model, images were split into 512 × 512 patches (∼ 300*µm* per side) and were subsequently encoded into 512-element feature vectors through CONCH. Models were trained using stochastic gradient descent (SGD) with a learning rate of 2×10^−5^ and a weight decay of 10^−5^. To address class imbalance between high-risk and low-risk images, a weighted sampling strategy was applied to balance class distributions within epochs. Same epoch schedule (50, 100, 150 epochs) was used to select the best-performing model by AUC.

For 2.5D analysis, neighboring levels at distances of *md* = 10, 20, 30, and 40 *µm* were ablated, with each range divided in different numbers of levels, *m* ∈ {1, 2, 3}. Aggregating a single pair of neighboring levels (*m* = 1) 10 *µm* apart from the target level yielded optimal performance.

#### Evaluation metrics

Model performance was primarily assessed using the Area Under the Receiver Operating Characteristic curve (AUC) for the ability to delineate between low- vs. high-risk images at a cohort level. To facilitate model selection during the development stage, we also computed balanced accuracy and F2 scores at 90% sensitivity for detecting high-risk samples. Compared to the F1 score, the F2 score emphasizes recall, which is important for a triage application (i.e. minimizing false negatives). It is calculated as:

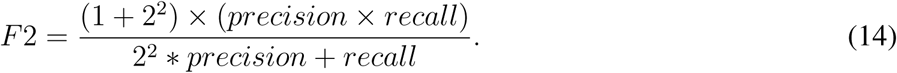

The 95% confidence intervals (CIs) were estimated via bootstrapping, generating *N* = 1000 bootstrap resamples by sampling with replacement for balanced accuracy and F2 score calculations ^85^. For AUC, accelerated Delong’s method was applied to derive the CIs.

Using the same bootstrapping strategies, statistical differences between methods are reported. Here we assume 2.5D models outperform their 2D counterparts. An empirical distribution of model performance difference (balanced accuracy and F2 score) was calculated at each resampled bootstrap (*N* = 1000) for the calculation of a one-side p-value, denoted as *p <* 0.05^∗^, *p <* 0.01^∗∗^, and *p <* 0.005^∗∗∗^. For AUC, accelerated Delong’s method was used to compare statistical differences between models.

#### Inference

For the purposes of evaluating the generalizability of models **Fig. 2d**, performing level-by-level predictions **Fig. 3**, as well as for triaging highest-risk levels for the clinical validation study **Fig. 4**, the final models were trained on all images within the development cohorts using the best hyperparameters identified during cross-validation.

### Model interpretability

#### Heatmap visualizations of patch attention

To generate fine-grained attention heatmaps, we extracted 2D patches with overlap to reduce blocky artifacts and computed corresponding attention scores, which were normalized to sum up to 1 within each level. An overlap of 50% was applied to prostate images and 75% to esophageal images. A coolwarm colormap, with red and blue colors indicating high and low attention values respectively, was then generated based on the normalized attention scores and overlaid on the raw false-colored images with a transparency value of 0.5.

#### Level attention

With CARP3D*_L_*_→*D*_, we computed level attention scores assigned by the depth-aggregation module to indicate the relative importance of the target level (**X_k_**) and its neighboring levels (**X_k_**_+**id**_).

### Level-by-level triage in 3D datasets

#### Risk-profile generation

During level-by-level triage, the final model (see inference section) was used to predict risk scores for 2D levels at 10 *µm* intervals within 3D pathology datasets, generating depth-wise risk profiles. These profiles were subsequently smoothed using a moving average, with a window size of 7 (corresponding to 60 *µm*) for prostate samples and a window size of 3 (20 *µm*) for esophagus samples.

#### PCA plot of highest-attended patches per level

For each level, the 15 patches that received the highest patch attention scores produced by **Eq. 1** were featurized using the CONCH encoder and projected along two dimensions via principal component analysis (PCA) as in **Fig. 3**.

#### Range of predicted risk across depth

To assess risk heterogeneity along the depth axis, we calculated the difference between the top 1 percentile and 99 percentile of predicted risk scores within each sample’s risk profile. In **Fig. 4b**, the swarm plot illustrates the range of predicted risk for individual samples, while the overlaid violin plot summarizes the overall distribution across the cohort.

### Preliminary clinical validation

#### Selection of AI-triaged levels

For prostate datasets, the three highest-risk levels were selected from each 3D volume based on risk profiles generated from level-by-level triage. To minimize the presentation of redundant information in different triaged levels, a minimum interval of 60 *µ*m was maintained between selected levels, based on discussions with genitourinary pathologists. For esophagus datasets, eight levels were selected with a minimum interval of 20 *µ*m between levels, as recommended by gastrointestinal pathologists. Triaged levels are selected in order of highest risk while maintaining the aforementioned minimum depth interval between triaged levels.

For esophagus datasets, we also compared diagnoses based on three AI-triaged levels versus eight AI-triaged levels in **Extended Data Fig. 8a**, separated by a one-month washout period. The results suggest that using eight levels can further improve the identification of dysplasia/cancer.

#### Comparison of standard H&E histology sections vs AI-triaged 3D pathology

To validate the clinical value of our methods, we compared diagnoses based on real H&E slides and AI-triaged 3D pathology images based on the same sample cohorts. After OTLS imaging, specimens were transferred into 100% EtOH and submitted for standard H&E histology, where 3 sections were cut for each prostate specimen at a spacing of 20 *µm* between levels (standard practice), and 16 consecutive sections were cut for each esophagus specimen (clinical practice at UWMC). A panel of five GU pathologists diagnosed prostate specimens based on 2D H&E sections as well as AI-triaged 3D pathology images, with a 1-month wash-out period between stages, while esophageal specimens were similarly diagnosed by three GI pathologists.

#### Agreement

We grouped samples into four categories based on the majority-opinion diagnosis for each modality: (1) low risk based on both standard 2D histology and AI-triaged 3D pathology, (2) high risk for both, (3) low risk for 2D histology but high risk for AI-triaged 3D pathology, and (4) high risk for 2D histology but low risk by AI-triaged 3D pathology. Within each group, we calculated the average percentage of pathologists who agreed with the assigned risk category, providing a measure of group-wise diagnostic consistency, as shown in **Fig. 4d,g**.

To further quantify the agreement between pathologists when intepreting 2D H&E histology sections and AI-triaged levels from 3D pathology, we reported Fleiss’ Kappa calculated on all samples in the cohort.

### Computational hardware and software

The computational work was performed on AMD Ryzen multicore CPUs (central processing units) with 256 GB of memory. Two NVIDIA GeForce RTX 3090 Ti GPUs (graphics processing units) were used for model training. The deep learning model implementation was based on Python 3.7 and PyTorch (version 1.7). CONCH (https://github.com/mahmoodlab/CONCH), pretrained with pathology image-text pairs, was used for patch feature extraction. Other Python libraries used for data processing and analysis include numpy (version 1.21), h5py (version 3.7), falsecolor-python (version 1.1.4), tifffile (version 2021.7.2), pandas (version 1.3.5), scipy (version 1.13.0), pillow (version 9.5.0), opencv-python (version 4.7.0), torchvision (version 0.8.0), timm (version 0.5.4), and matplotlib (version 3.5.3).

## Data and code availability

CARP3D code will be made available upon publication. We will release all 3D pathology images used for prostate and esophagus model development and their corresponding annotations for research and educational use.

## Author contributions

G. Gao, A.H. Song, L.A.E. Barner, W.M. Grady, F. Mahmood, and J.T.C. Liu conceived the study and designed the experiments. P. Lal, A. Madabbushi, W. Burke, L.D. True, W.M. Grady and J.T.C. Liu collected clinical samples. G. Gao, L.A.E. Barner, F. Wang, D. Brenes, S.S.L. Chow, R. Wang, and K.W. Bishop stained, imaged and processed samples for the 3D pathology data cohorts. Y. Liu, X. Farre, M. Divatia, M.R. Downes, F. Vakar-Lopez, P. Lal, L.D. True, and D.M. Reddi annotated the 3D pathology data and performed the clinical validation studies. G. Gao and R. Yan ran all the computational experiments with guidance from A.H. Song. G. Gao, A.H. Song, and H. Hsieh analyzed the clinical validation results. G. Gao, R. Yan, A.H. Song, H. Hsieh, and J.T.C. Liu prepared the manuscript. H. Hsieh designed the video abstract. All authors contributed to the writing. J.T.C. Liu supervised the research.

## Acknowledgements

This research was primarily supported by the National Institute of Diabetes and Digestive and Kidney Diseases (NIDDK) under R01DK138948 (Liu, Mahmood, and Grady) and the National Cancer Institute (NCI) under R01CA268207 (Liu and Madabhushi). NIH support for WM Grady includes U01CA152756, R01CA220004, U2CCA271902, U54CA163060, and U01CA182940. Funding for WM Grady is also provided by the Prevent Cancer Foundation, Cottrell Family Fund, Evergreen Fund, and Listwin Foundation to WMG. These studies are also supported by GiCaRes from UW Departments of Medicine (Gastroenterology Division) and Lab Medicine & Pathology. Additional NIH support includes grants R01EB031002 (Liu), R01CA268287 (Madabhushi), U01CA269181(Madabhushi), R01CA249992 (Madabhushi), R01CA202752 (Madabhushi), R01CA208236 (Madabhushi), R01CA216579 (Madabhushi), R01CA220581 (Madabhushi), R01CA257612 (Madabhushi), 1U01CA239055 (Madabhushi), 1U01CA248226 (Madabhushi), 1U54CA254566 (Madabhushi), 1R01HL15127701A1 (Madabhushi) and R01HL15807101A1 (Madabhushi). Support was also received from the VA Merit Review Award IBX004121A (Madabhushi) and sponsored research agreements from Bristol Myers-Squibb, and As-trazeneca. Additional support was through the Department of Defense (DoD) Prostate Cancer Research Program (PCRP) grant W81XWH-18-10358 (Liu and True), W81XWH-14-2-0183 (True), W81XWH-20-1-0851 (Madabhushi and Liu), the Pacific Northwest Prostate Cancer SPORE P50CA97186 (True), NSF Graduate Research Fellowship DGE-1762114 (K.W.B.), and the Canary Foundation. Research was also supported in part by the ARPA-H under contract number D24AC00357 (Liu) and D25AC00140 (Madabhushi). The content is solely the responsibility of the authors and does not necessarily represent the official views of the National Institutes of Health, the U.S. Department of Veterans Affairs, the Department of Defense, or the United States Government.

---

**Extended Data Figure 1:**
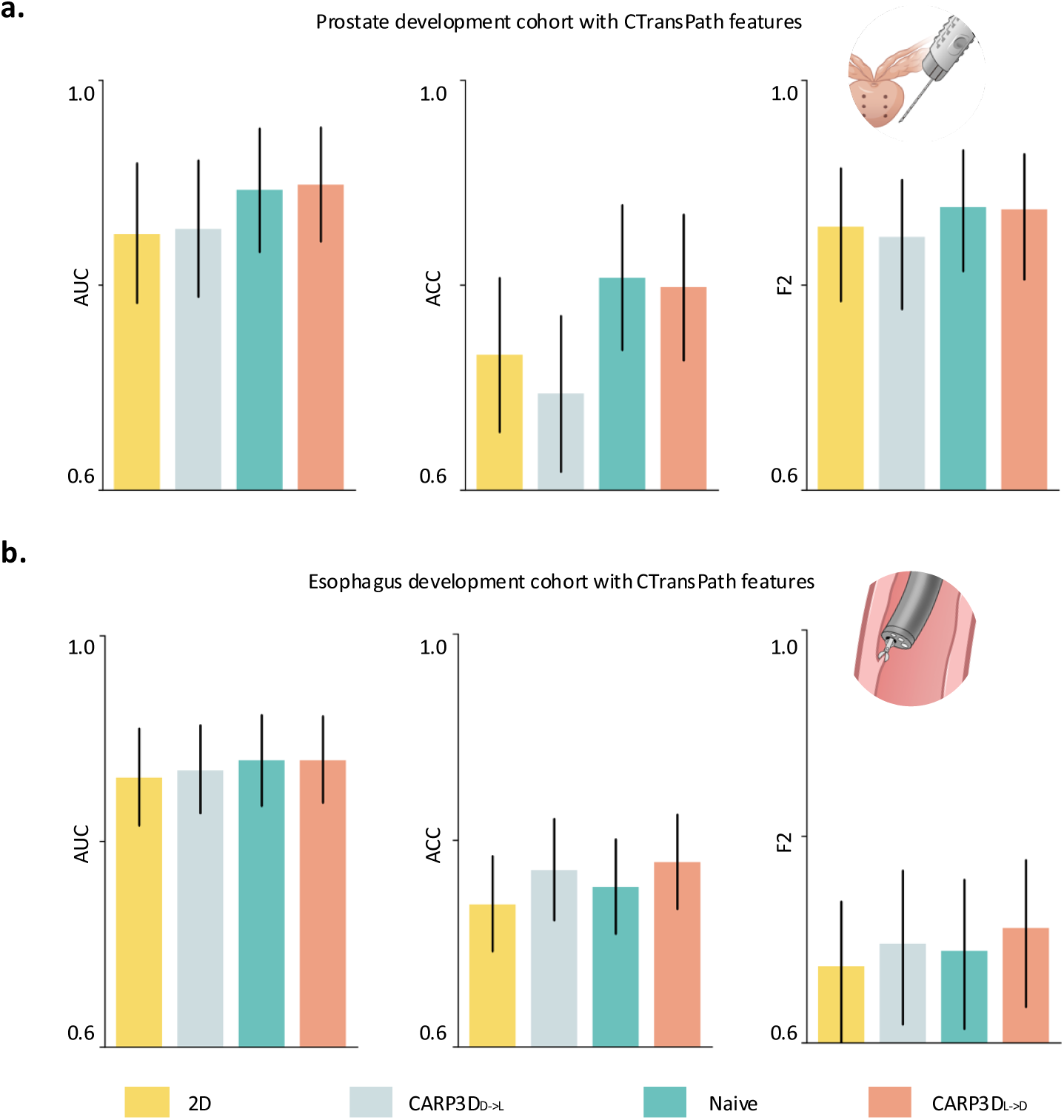
Cohort-level metrics comparing different aggregation strategies based on CTransPath patch features. AUC, balanced accuracy, and F2 scores for 2D, CARP3D*_D_*_→_*_L_*, CARP3D*_N_*, and CARP3D*_L_*_→_*_D_* aggregation strategies using CTransPath features on the **(a)** prostate and **(b)** esophagus development cohort, based on leave-one-out cross-validation. CTransPath patch feature encoder ^42^ is a Swin Transformer pretrained on a combination of 33K WSIs from TCGA and the Pathology AI Platform (PAIP).

**Extended Data Figure 2:**
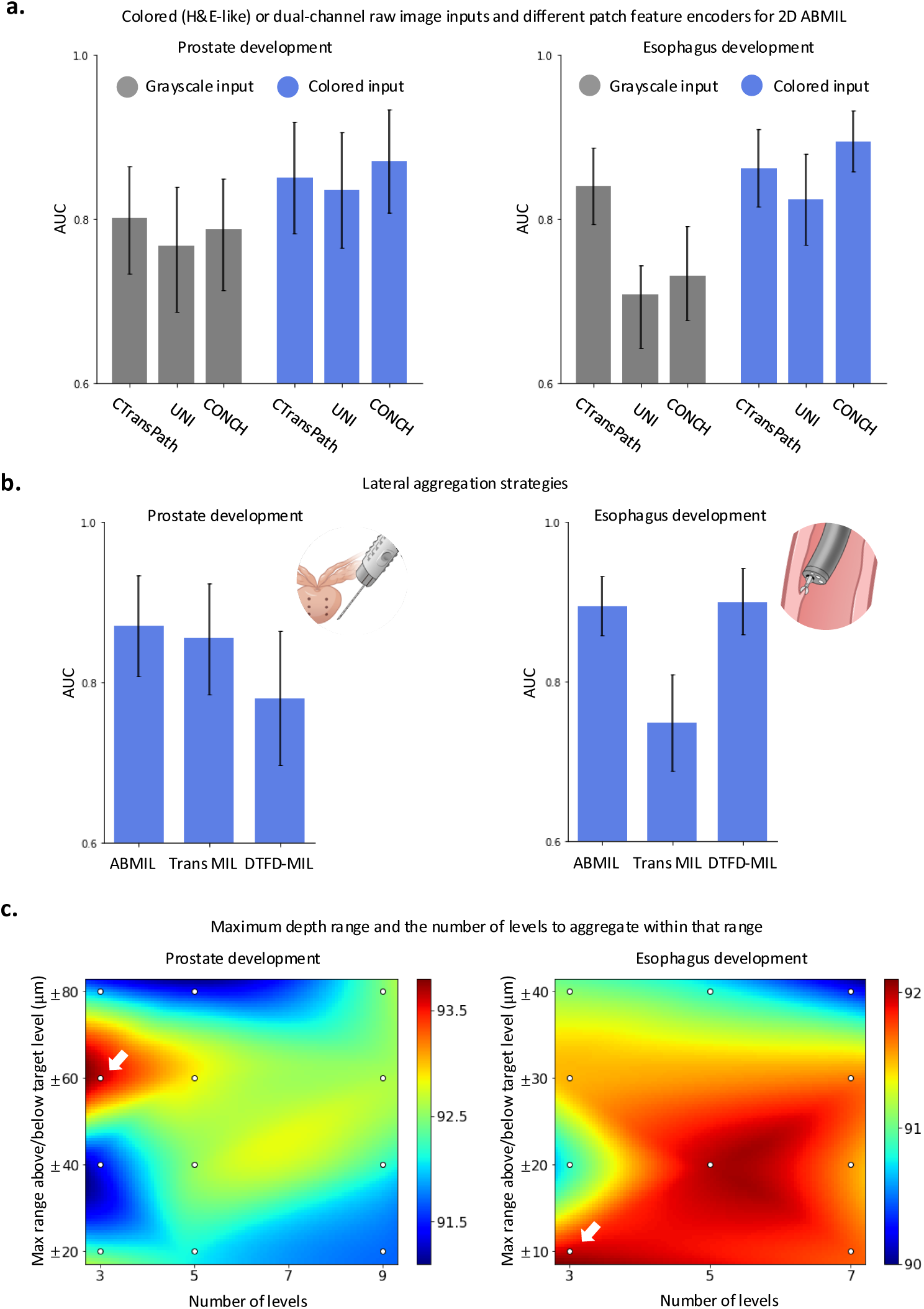
Optimization of key design parameters for CARP3D. First, we optimize 2D MIL model design by ablating the input format, patch feature encoder, and lateral aggregation strategies. Then, using the same optimized design, we tune the maximum depth range and the number of levels aggregated with CARP3D. **(a)** Ablation analysis for false-colored (H&E-like) or dual-channel inputs (RGB-format with nuclei in the red channel, eosin in green, and blank blue channel) and patch feature encoders (CONCH ^71^, UNI ^45^, CTransPath ^42^) for 2D ABMIL. The results show that self-supervised models (trained on H&E whole slide images) perform best with H&E-false-colored inputs, with CONCH achieving the best performance. **(b)** Ablation analysis for lateral aggregation strategies (ABMIL ^34^, TransMIL ^38^, DTFD-MIL ^39^), with ABMIL showing superior results. **(c)** Ablation analysis for the maximum depth range being aggregated (y axis) and the number of levels to aggregate within that range (x axis). The heatmap is interpolated from discrete evaluation points (white dots), each corresponding to a combination of depth range and number of levels. The optimal configuration chosen in our study is indicated by the white arrow.

**Extended Data Figure 3:**
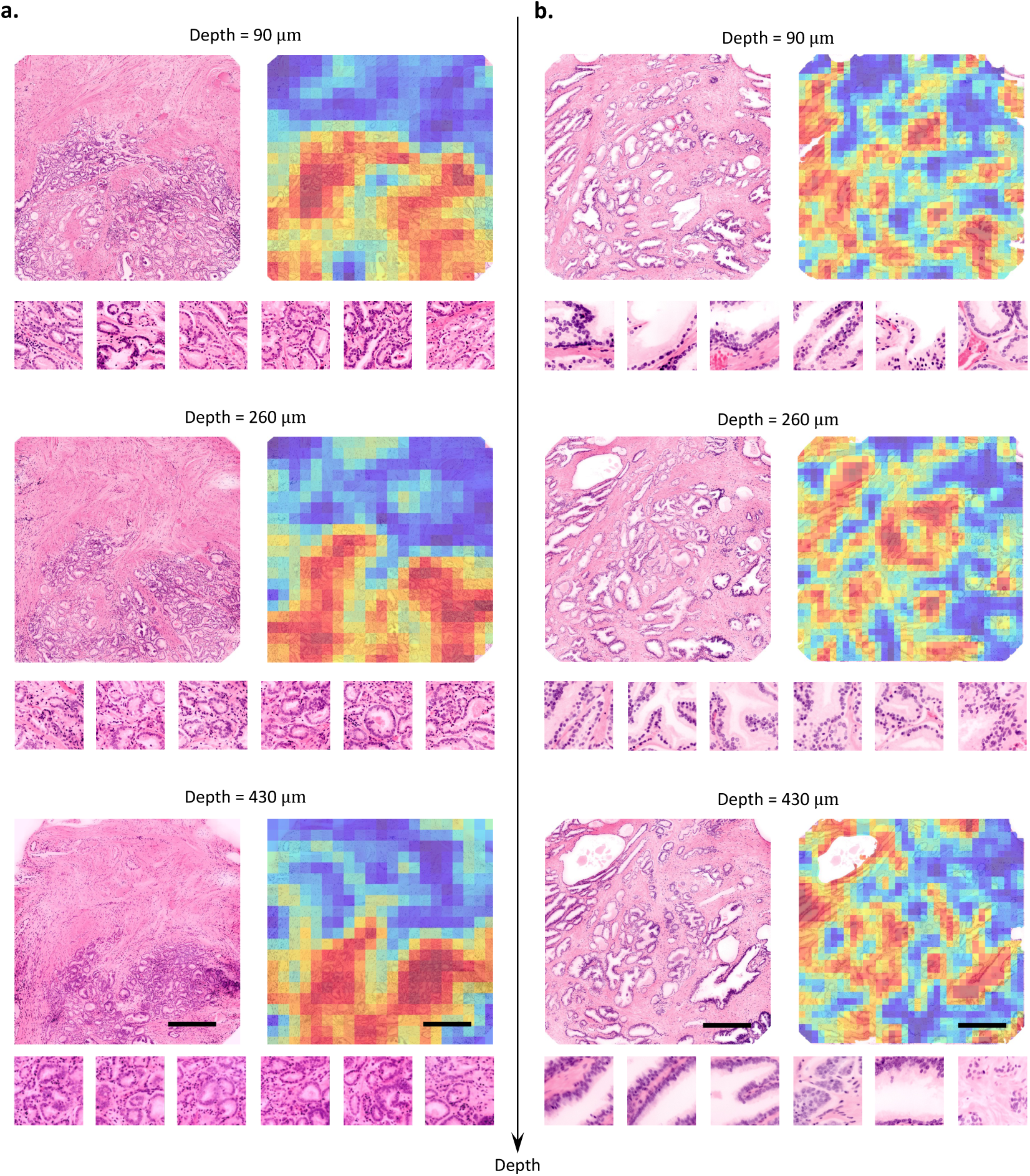
Examples of patch attention heatmaps for the prostate cohort. **(a)** Higher-grade prostate cancer. **(b)** Low-grade prostate cancer. Attention heatmaps are shown for three representative levels at different depths within the 3D datasets, with 50% overlap between patches. High attention is focused on glandular structures in both **(a)** and **(b)**, with **(a)** showing crowded and fused glands (Gleason pattern 4) and **(b)** showing discrete cancer glands (Gleason pattern 3) and large benign glands. Scale bars are 500 *µm* and patches sizes are ∼ 200 *µm*.

**Extended Data Figure 4:**
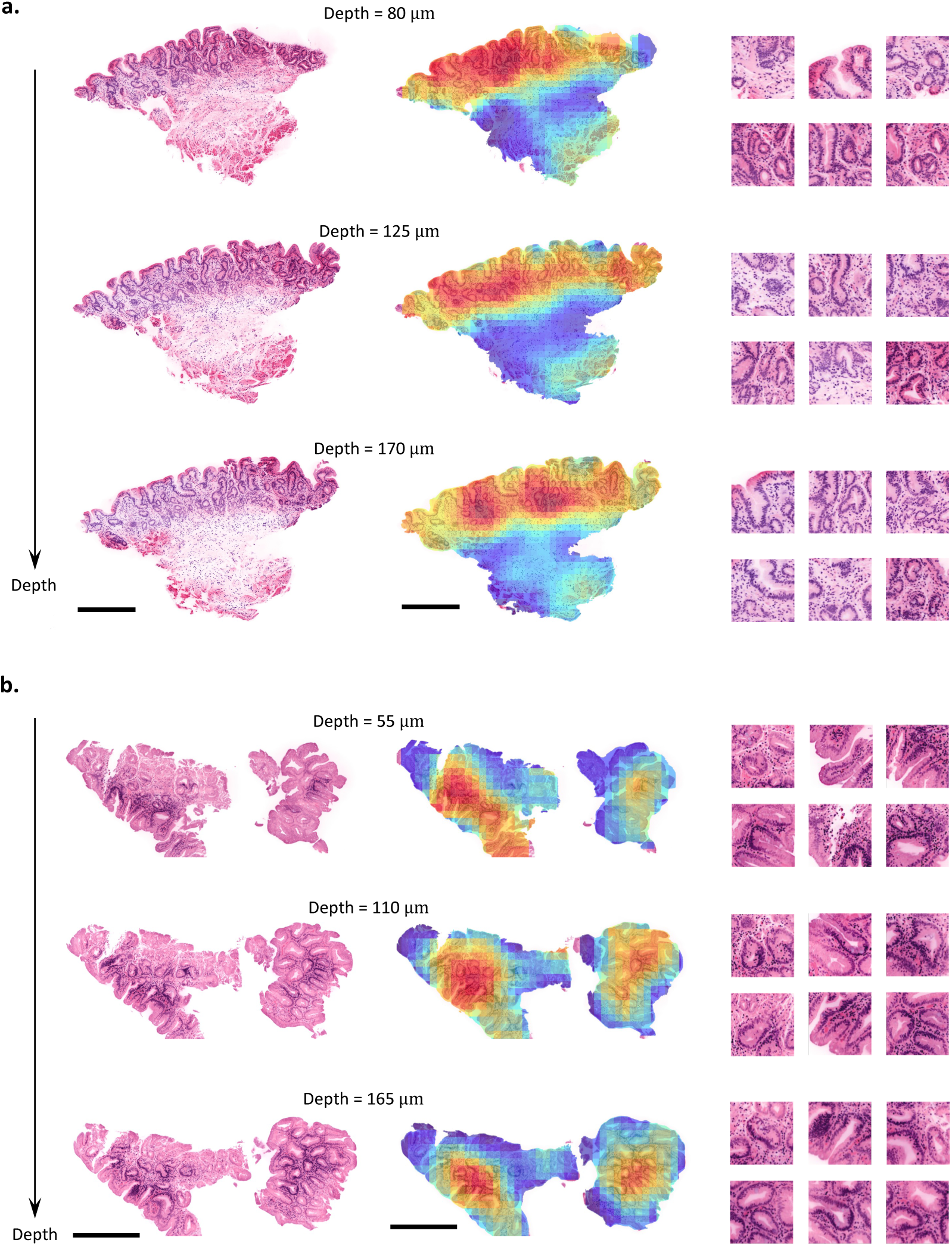
Examples of patch attention heatmaps for the esophagus cohort. **(a)** Dysplasia/cancer. **(b)** Benign. Attention heatmaps are shown for three representative levels at different depths within the 3D datasets, with 75% overlap between patches. High attention is focused on epithelial surface regions in both **(a)** and **(b)**, with **(a)** showing glands characterized by dark elongated nuclei with a stratified arrangement and prominent mitoses (low-grade dysplasia), and **(b)** showing glands with goblet cells and mucin caps indicative of a mature surface (nondysplastic Barrett’s esophagus). Scale bars are 500 *µm* and patches sizes are 200 *µm*.

**Extended Data Figure 5:**
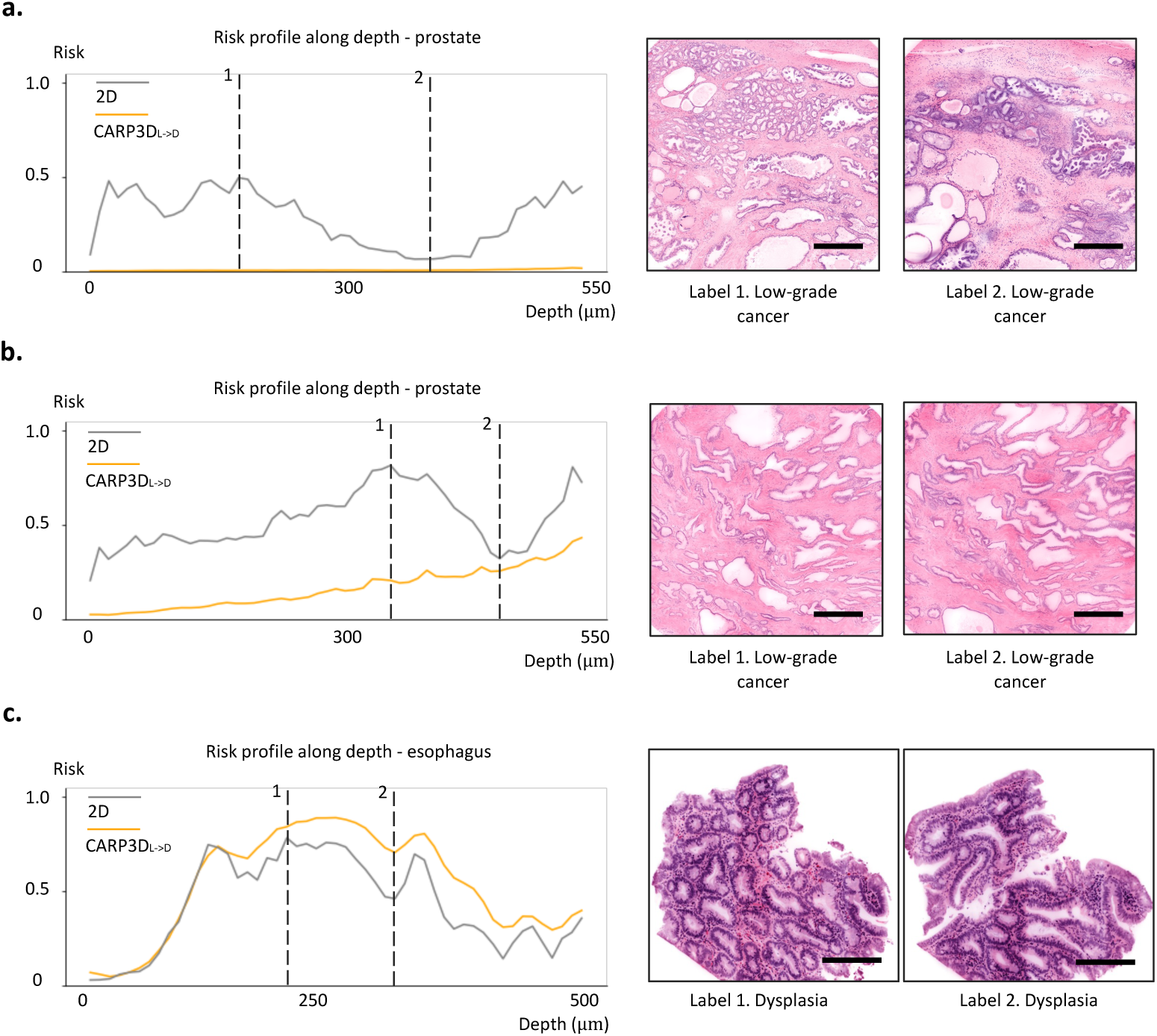
Examples of predicted risk profiles as a function of depth for the prostate and esophageal cohorts. Risk profiles for three representative datasets with CARP3D_L→D_ vs. 2D aggregation are shown (grey: 2D, orange: CARP3D_L→D_). The results demonstrate that CARP3D_L→D_ produces more stable and accurate predictions, while 2D aggregation results in noisier results. This is supported by the pathologist-assigned labels aligning more closely with CARP3D_L→D_ predictions for example levels. **(a)**, **(b)** Low-grade prostate cancer across depth. **(c)** Dysplasia identified throughout the central region. Scale bars are 500 *µm*.

**Extended Data Figure 6:**
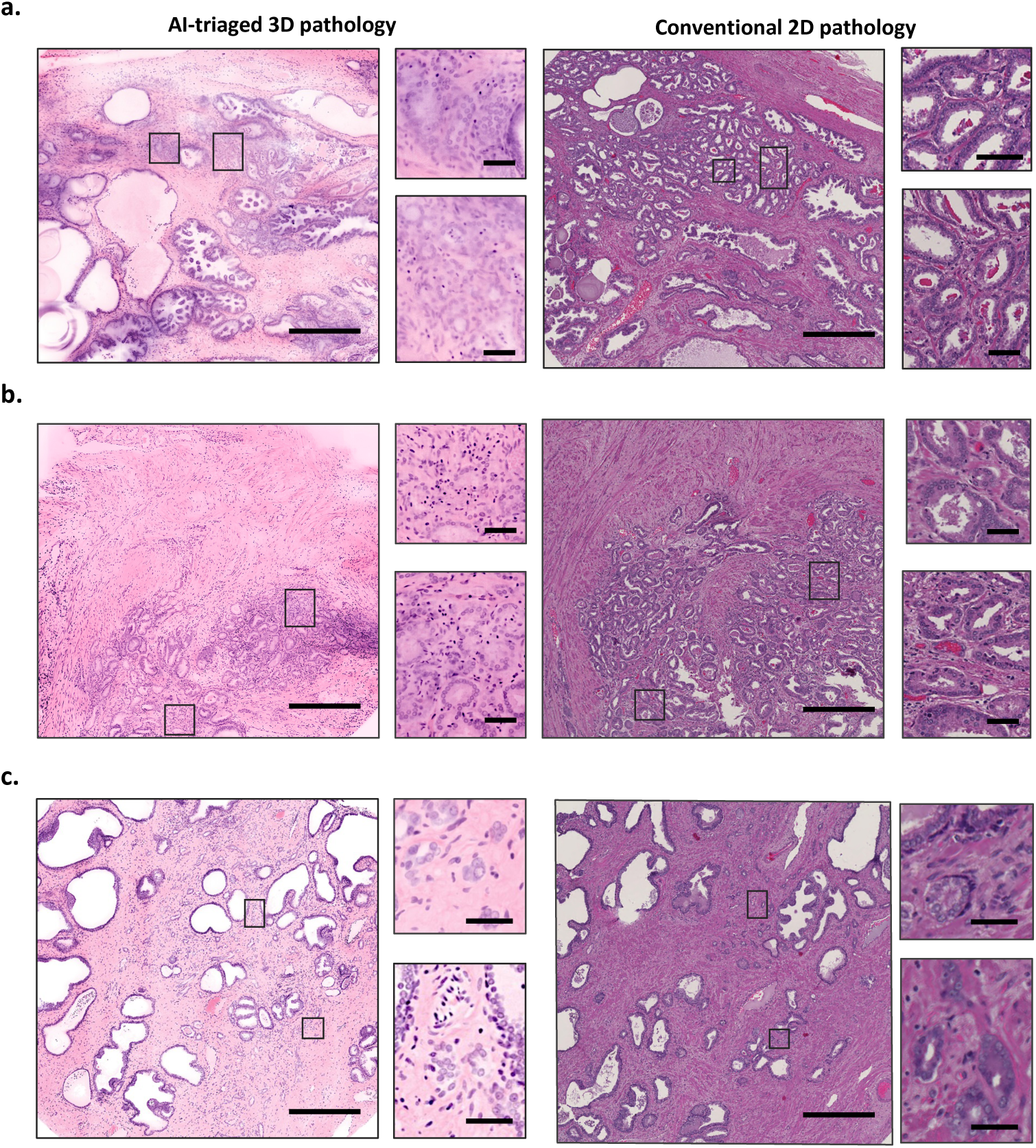
Additional examples of upgraded cases from Fig. 4d, comparing AI-triaged 3D pathology (left) with standard 2D histopathology (right). **(a, b)** Fused glands (Gleason pattern 4) are identified on the left (AI-triaged) levels, while conventional 2D histopathology only shows distinct glands (Gleason pattern 3). **(c)** Poorly-formed glands are identified on the left, while conventional 2D histopathology only shows fully formed glands with clear lumens (Gleason pattern 3). Scale bars are 500 *µm* for large images and 100 *µm* for zoomed-in regions.

**Extended Data Figure 7:**
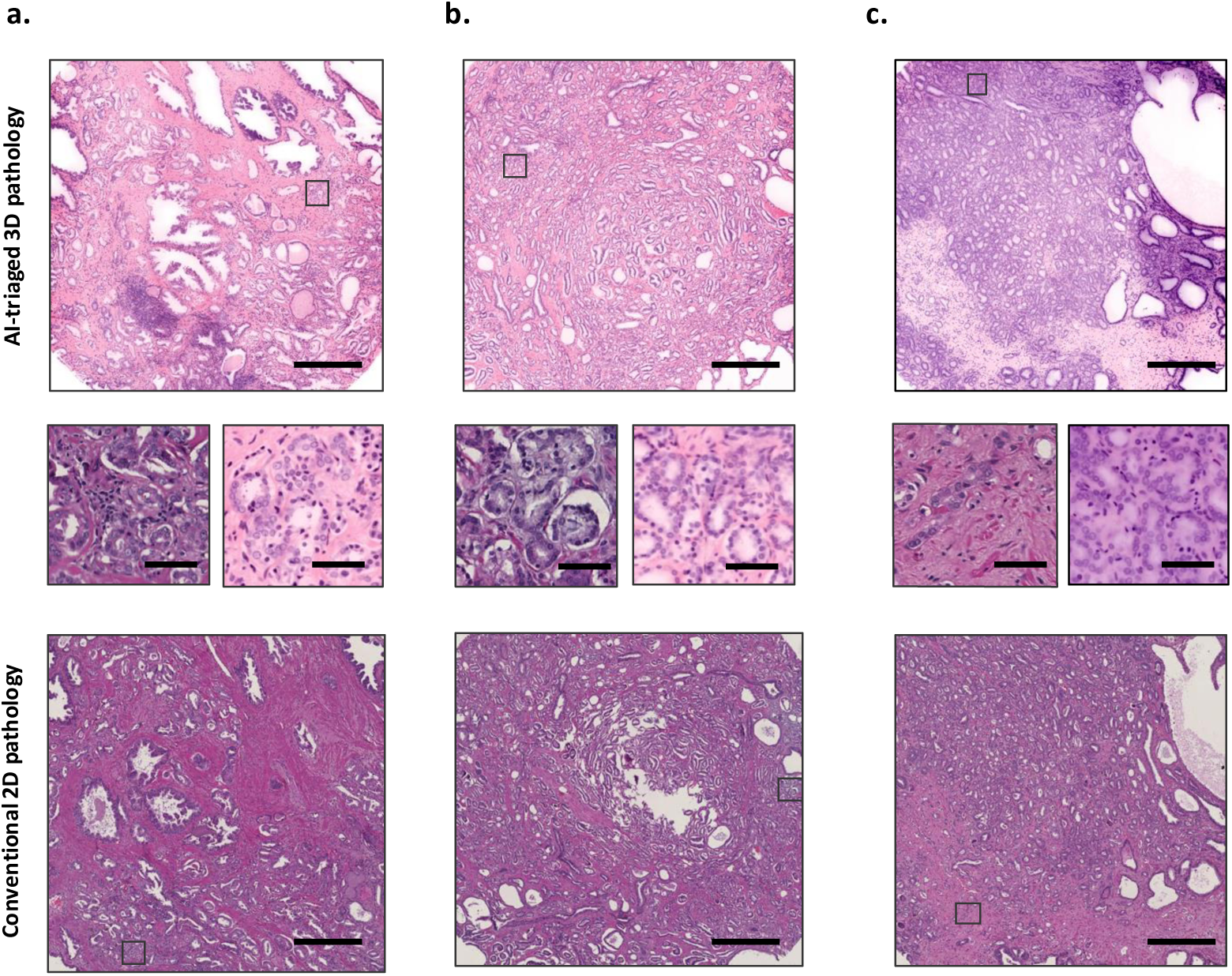
Downgraded cases with AI-triaged 3D pathology for the prostate cancer cohort. Both the AI-triaged levels and 2D slides contain only small foci that are suspicious of containing higher-grade prostate cancer, contributing to low agreement with the downgrading categorization in Fig. 4d. For the three cases, **(a)-(c)**, two pathologists determined that neither AI-triaged images nor arbitrary 2D sections contained higher-grade prostate cancer. One pathologist labeled both 2D and AI-triaged images as containing higher-grade prostate cancer, while two others identified higher-grade cancer only on the 2D slides. Scale bars are 500 *µm* for large images and 100 *µm* for zoomed-in regions.

**Extended Data Figure 8:**
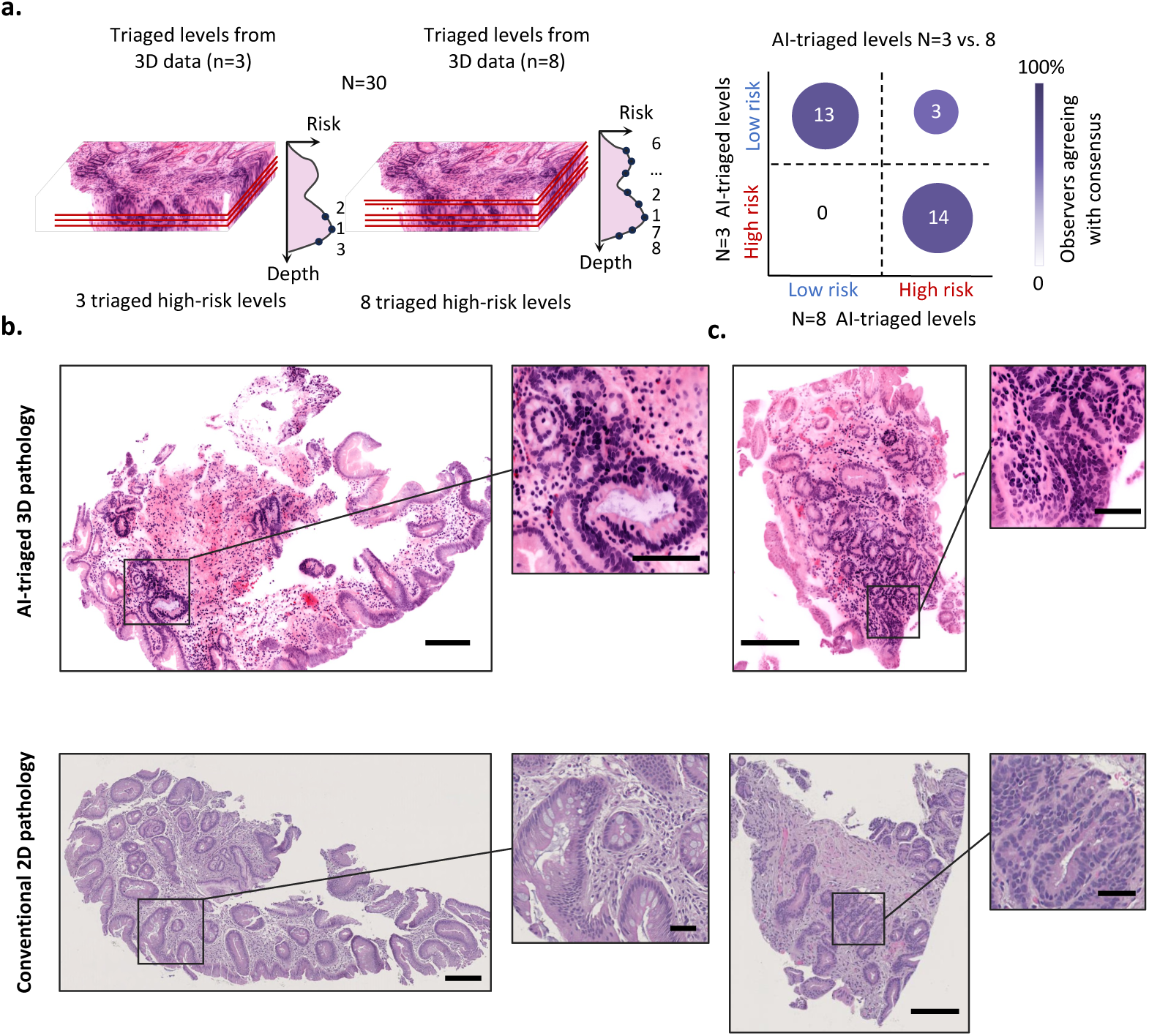
Clinical validation with the esophagus cohort. **(a)** Diagnosis comparisons based on three AI-triaged levels vs. eight AI-triaged levels with the same panel of pathologists and sample cohort. The examination of eight levels by pathologists enabled the identification of three additional high-risk samples that were missed when only three levels were examined. **(b, c)** Additional examples of upgraded cases from Fig. 4g., comparing AI-triaged 3D pathology (top) with conventional 2D pathology (bottom). Dysplasia was identified by AI-triaged 3D pathology, whereas only Barrett’s esophagus was seen with 2D histopathology. Scale bars are 500 *µm* for large images and 100 *µm* for zoomed-in regions.

**Extended Data Table 1:**
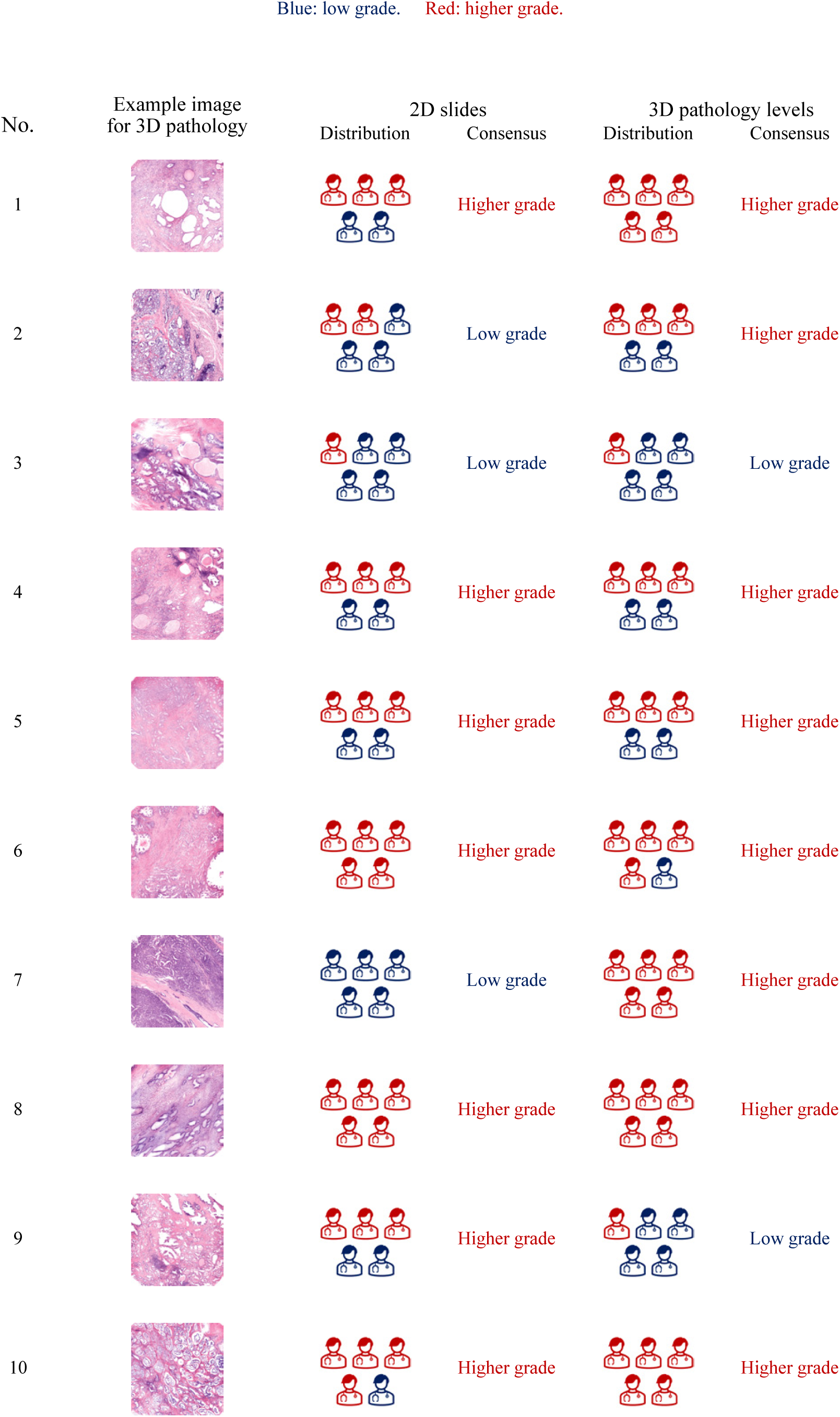
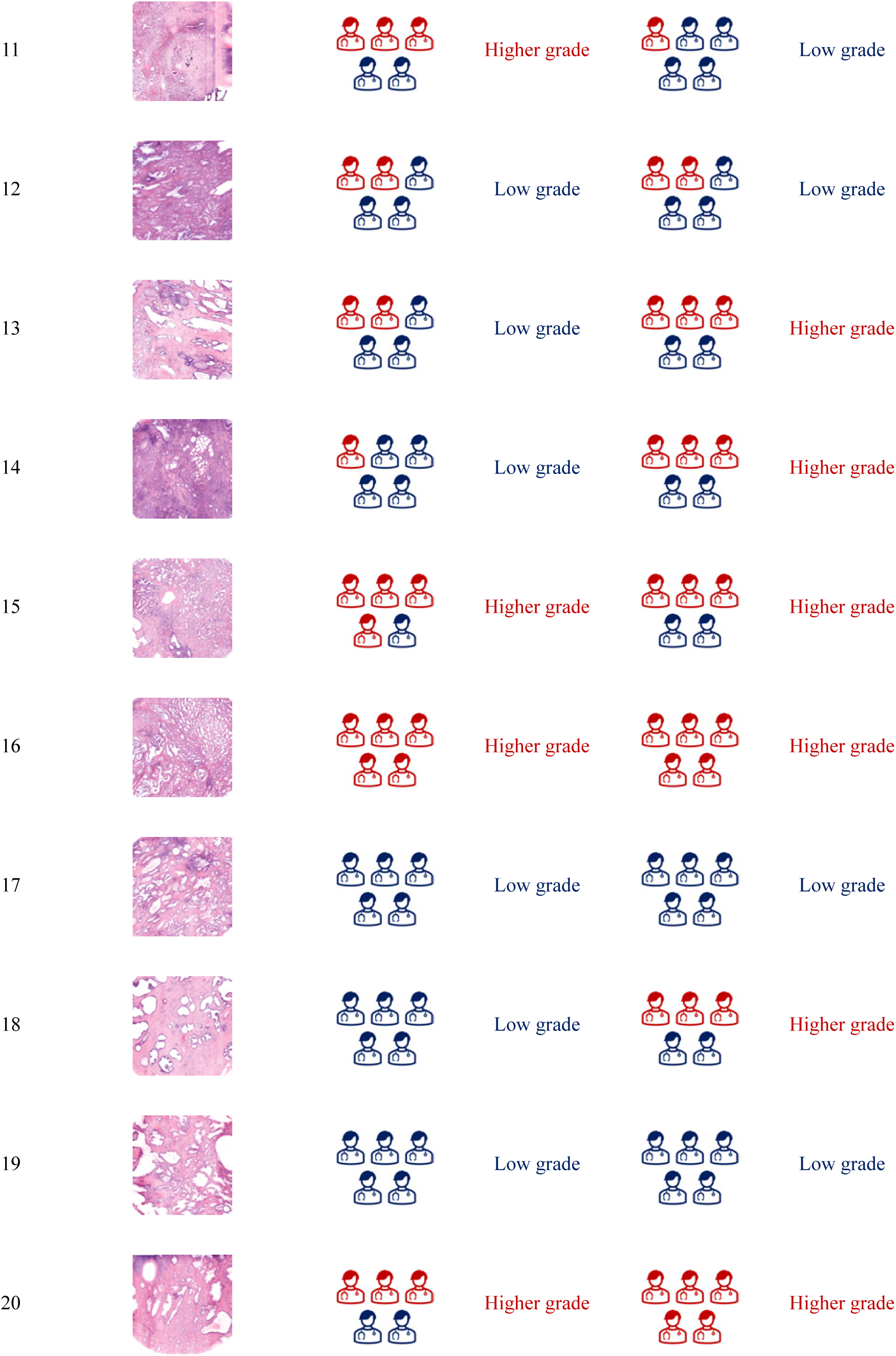
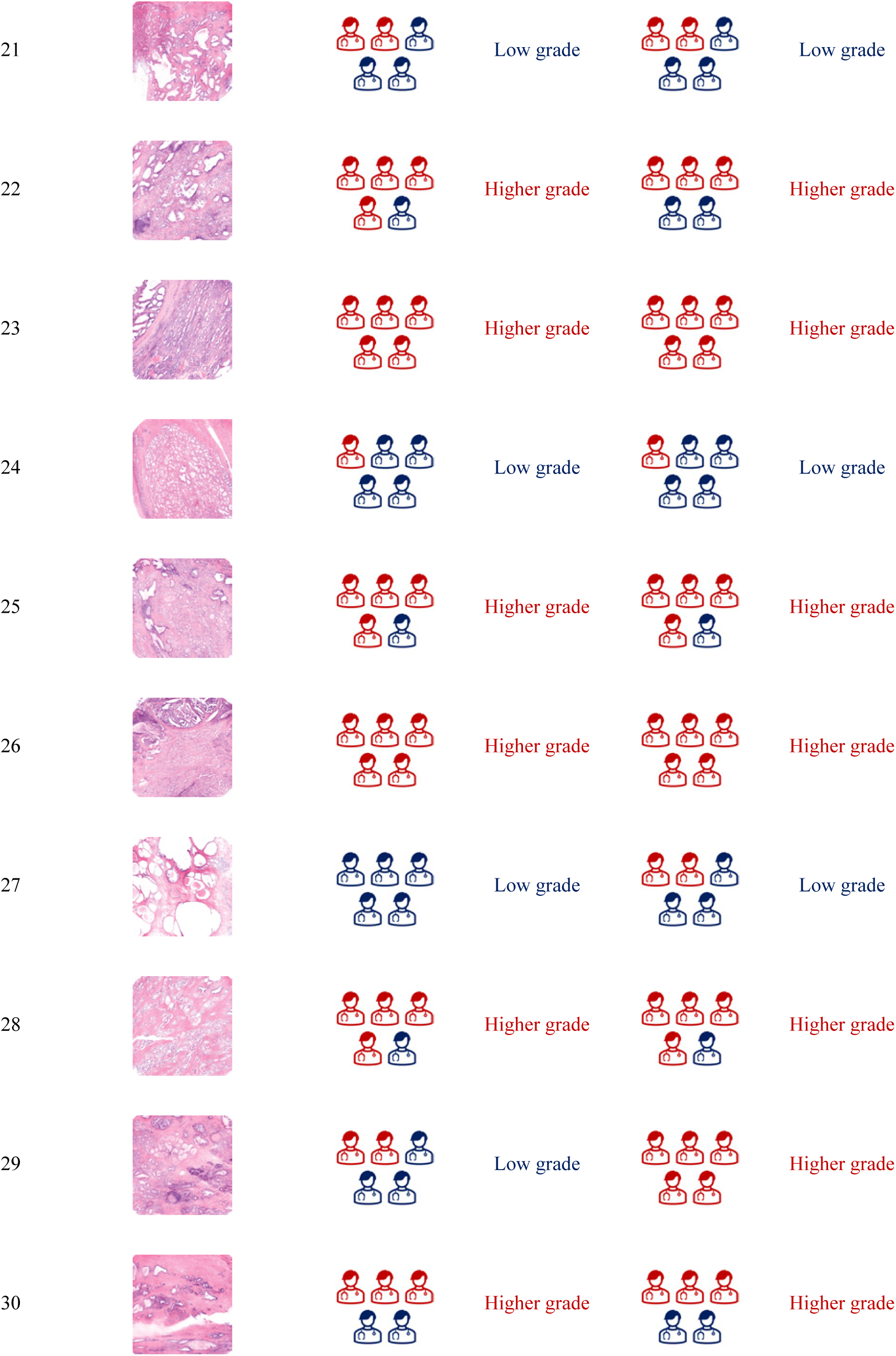
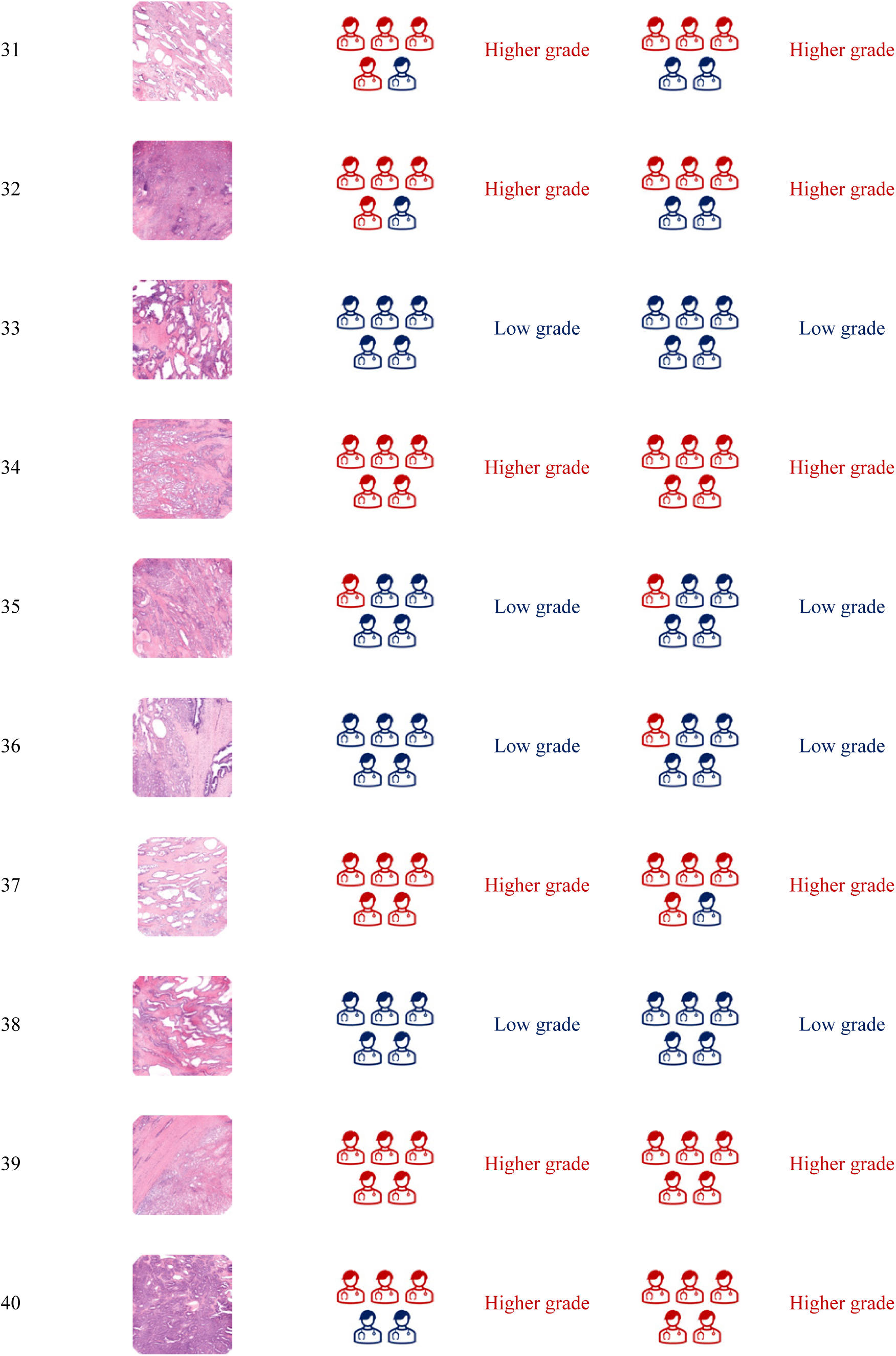
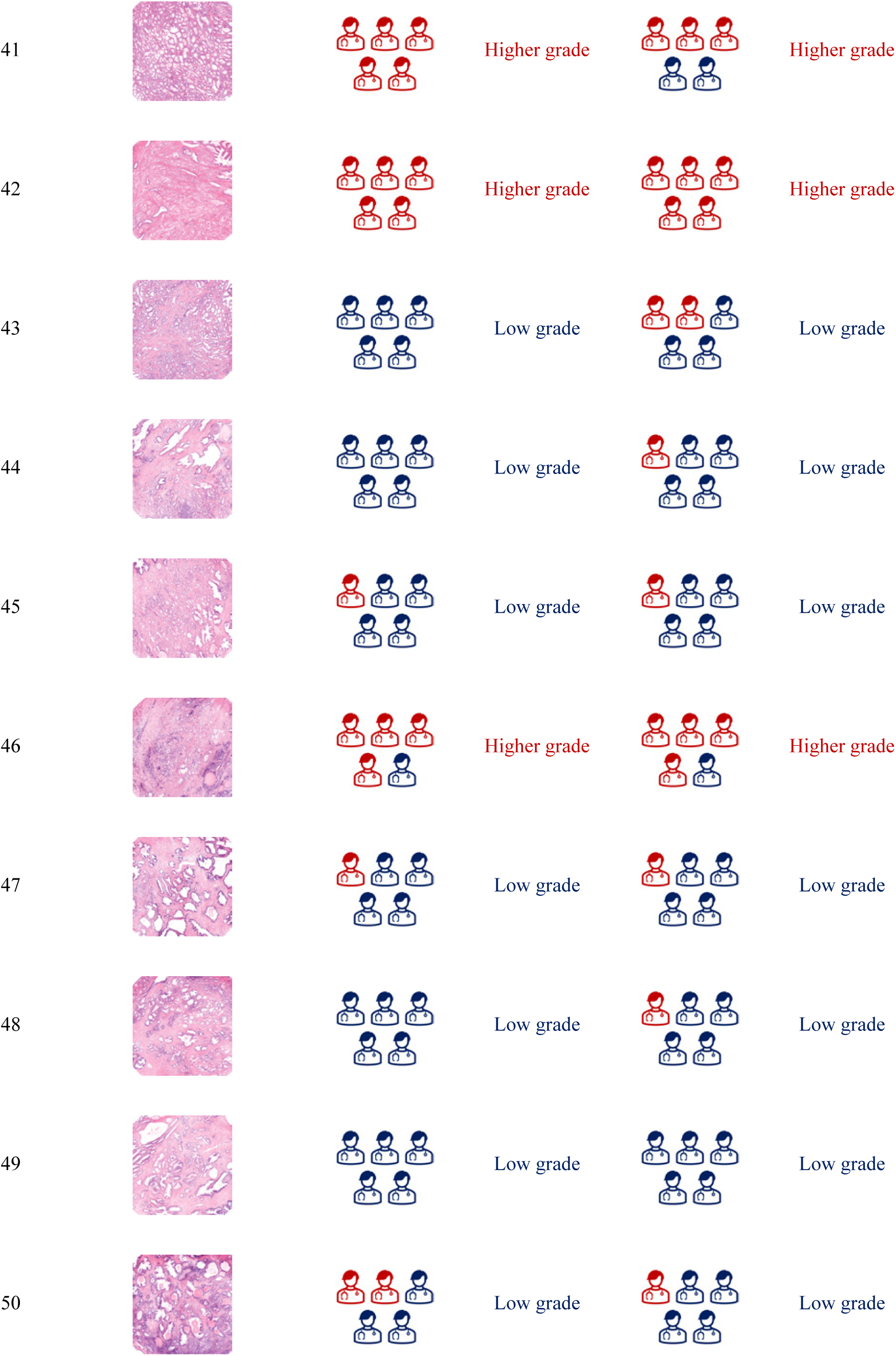
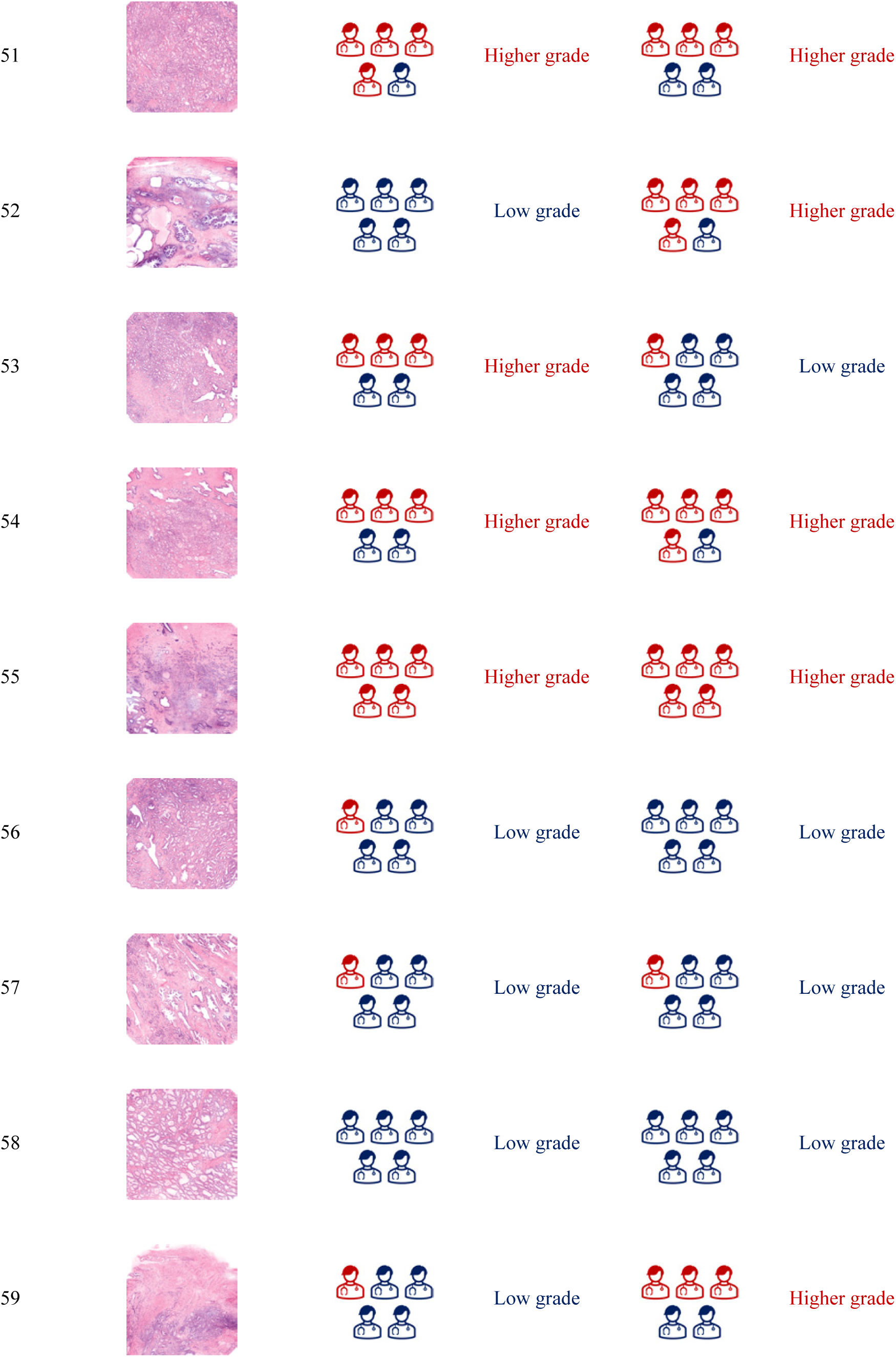
Dataset for CARP3D prostate clinical validation.

**Extended Data Table 2:**
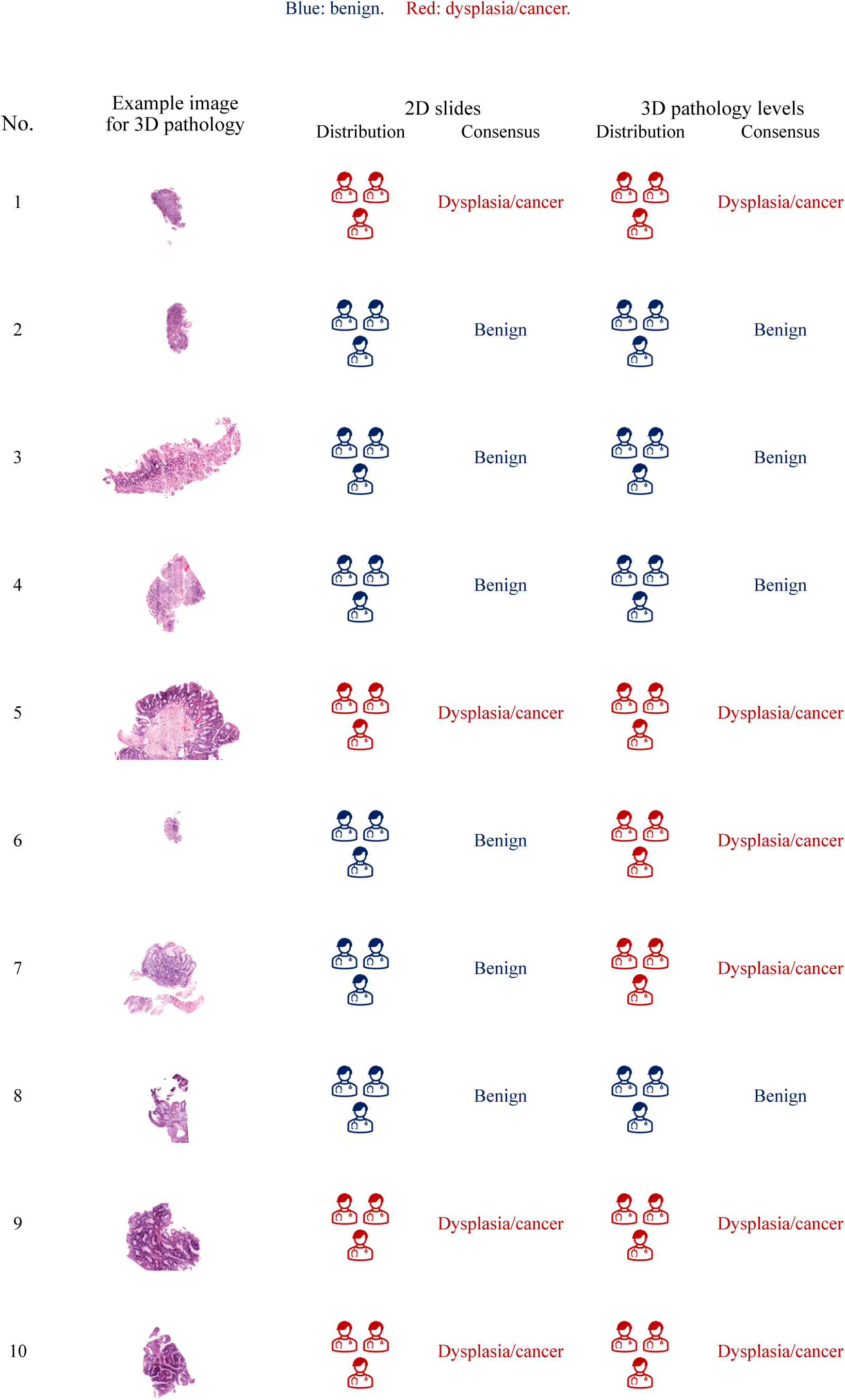
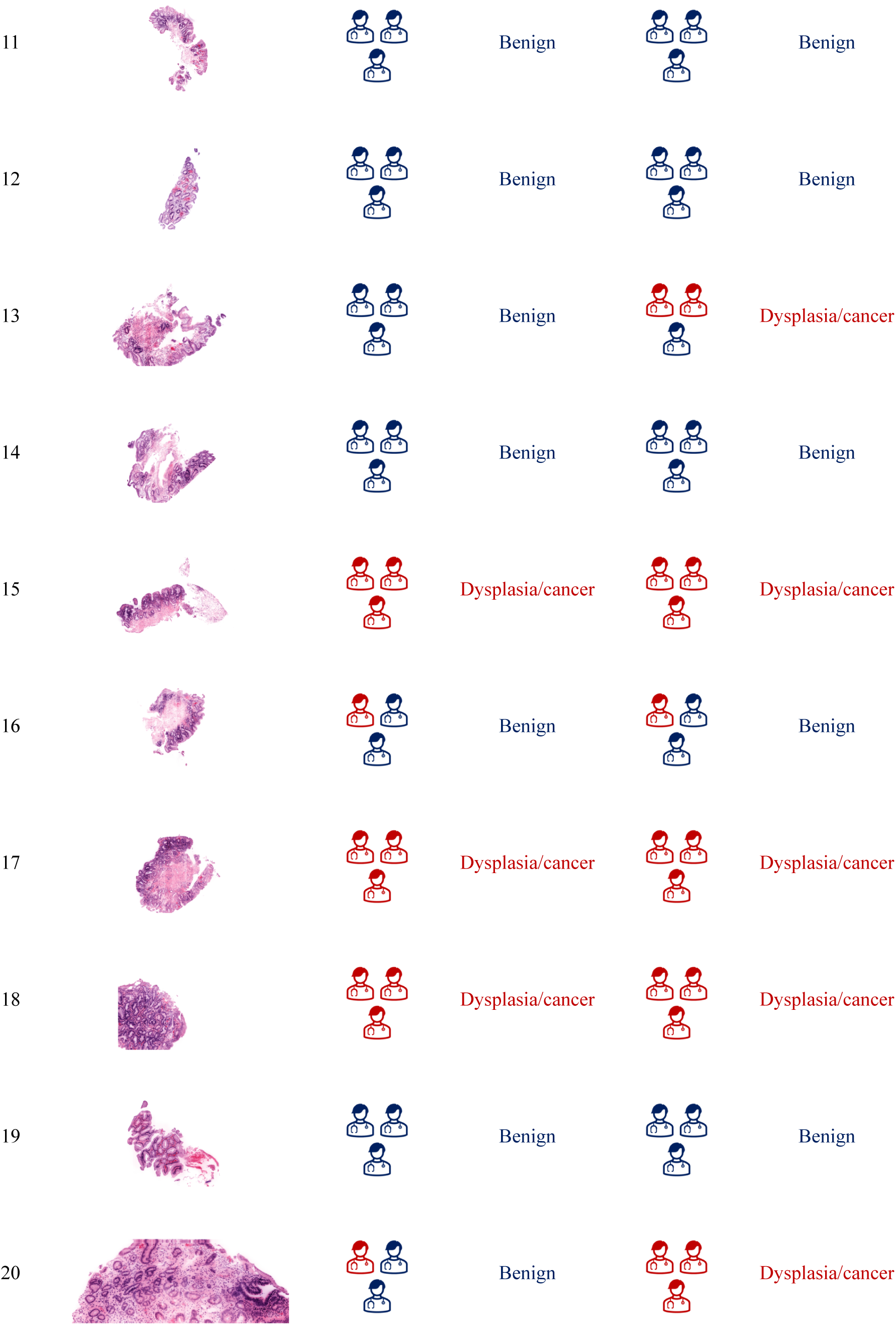
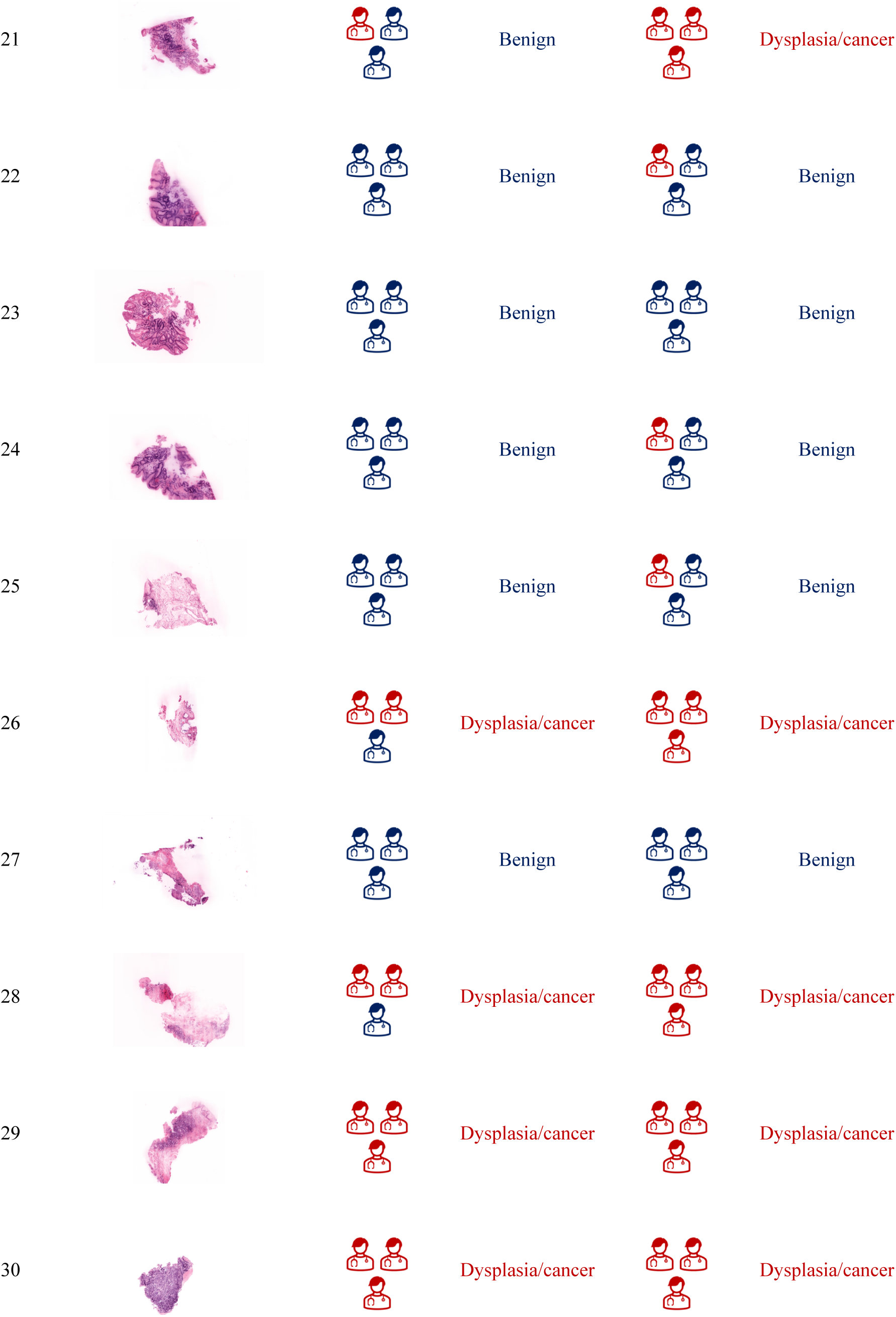
Dataset for CARP3D esophagus clinical validation.

## References

1. Liu, J. T. et al. Harnessing non-destructive 3D pathology. Nature biomedical engineering 5, 203–218 (2021).

2. Liu, J. T. et al. Engineering the future of 3D pathology. The Journal of Pathology: Clinical Research 10, e347 (2024).

3. La, E. B., et al. AI-triaged 3D pathology to improve detection of esophageal neoplasia while reducing pathologist workloads. Modern Pathology 100322–100322 (2023).

4. Koyuncu, C. et al. Visual assessment of 2-dimensional levels within 3-dimensional pathology data sets of prostate needle biopsies reveals substantial spatial heterogeneity. Laboratory Investigation 103, 100265 (2023).

5. Reddi, D. M. et al. Nondestructive 3D pathology image atlas of Barrett esophagus with open-top light-sheet microscopy. Archives of Pathology & Laboratory Medicine 147, 1164–1171 (2023).

6. Chitkara, Y. K. & Eyre, C. L. Evaluation of initial and deeper sections of esophageal biopsy specimens for detection of intestinal metaplasia. American journal of clinical pathology 123, 886–888 (2005).

7. Reyes, A. O. & Humphrey, P. A. Diagnostic effect of complete histologic sampling of prostate needle biopsy specimens. American journal of clinical pathology 109, 416–422 (1998).

8. Arista-Nasr, J. et al. Atypical small acinar proliferation: utility of additional sections and immunohistochemical analysis of prostatic needle biopsies. Nephro-urology monthly 4, 443 (2012).

9. Kiemen, A. L. et al. CODA: quantitative 3D reconstruction of large tissues at cellular resolution. Nature Methods 19, 1490–1499 (2022).

10. Lin, J.-R. et al. Multiplexed 3D atlas of state transitions and immune interaction in colorectal cancer. Cell 186, 363–381 (2023).

11. Forjaz, A. et al. Three-dimensional assessments are necessary to determine the true, spatially resolved composition of tissues. Cell Reports Methods 5, 101075 (2025).

12. Song, A. H. et al. Analysis of 3D pathology samples using weakly supervised AI. Cell 187, 2502–2520 (2024).

13. Bishop, K. W. et al. An end-to-end workflow for nondestructive 3D pathology. Nature Protocols 19, 1122–1148 (2024).

14. Barner, L. A., Glaser, A. K., Huang, H., True, L. D. & Liu, J. T. Multi-resolution open-top light-sheet microscopy to enable efficient 3d pathology workflows. Biomedical Optics Express 11, 6605–6619 (2020).

15. Glaser, A. K. et al. Light-sheet microscopy for slide-free non-destructive pathology of large clinical specimens. Nature biomedical engineering 1, 0084 (2017).

16. Glaser, A. K. et al. Multi-immersion open-top light-sheet microscope for high-throughput imaging of cleared tissues. Nature communications 10, 2781 (2019).

17. Glaser, A. K. et al. A hybrid open-top light-sheet microscope for versatile multi-scale imaging of cleared tissues. Nature methods 19, 613–619 (2022).

18. Bishop, K. W. et al. Axially swept open-top light-sheet microscopy for densely labeled clinical specimens. Optics Letters 49, 3794–3797 (2024).

19. Tang, R. et al. Micro-computed tomography (Micro-CT): a novel approach for intraoperative breast cancer specimen imaging. Breast cancer research and treatment 139, 311–316 (2013).

20. van Royen, M. E. et al. Three-dimensional microscopic analysis of clinical prostate specimens. Histopathology 69, 985–992 (2016).

21. Susaki, E. A. et al. Advanced cubic protocols for whole-brain and whole-body clearing and imaging. Nature protocols 10, 1709–1727 (2015).

22. Lee, M. Y. et al. Fluorescent labeling of abundant reactive entities (flare) for cleared-tissue and super-resolution microscopy. Nature protocols 17, 819–846 (2022).

23. Matsumoto, K. et al. Advanced cubic tissue clearing for whole-organ cell profiling. Nature protocols 14, 3506–3537 (2019).

24. Ertürk, A., et al. Three-dimensional imaging of solvent-cleared organs using 3DISCO. Nature protocols 7, 1983–1995 (2012).

25. Richardson, D. S. et al. Tissue clearing. Nature Reviews Methods Primers 1, 84 (2021).

26. Hörl, D., et al. BigStitcher: reconstructing high-resolution image datasets of cleared and expanded samples. Nature methods 16, 870–874 (2019).

27. Serafin, R., Xie, W., Glaser, A. K. & Liu, J. T. Falsecolor-python: a rapid intensity-leveling and digital-staining package for fluorescence-based slide-free digital pathology. Plos one 15, e0233198 (2020).

28. Braxton, A. M. et al. 3D genomic mapping reveals multifocality of human pancreatic precancers. Nature 629, 679–687 (2024).

29. Joshi, S. et al. InterpolAI: deep learning-based optical flow interpolation and restoration of biomedical images for improved 3D tissue mapping. Nature Methods 1–12 (2025).

30. Xie, W. et al. Prostate cancer risk stratification via nondestructive 3d pathology with deep learning–assisted gland analysis. Cancer research 82, 334–345 (2022).

31. Serafin, R. et al. Nondestructive 3D pathology with analysis of nuclear features for prostate cancer risk assessment. The Journal of pathology 260, 390–401 (2023).

32. Gao, G. et al. Triage of 3D pathology data via 2.5D multiple-instance learning to guide pathologist assessments. In Proceedings of the IEEE/CVF Conference on Computer Vision and Pattern Recognition, 6955–6965 (2024).

33. Song, A. H. et al. Artificial intelligence for digital and computational pathology. Nature Reviews Bioengineering 1, 930–949 (2023).

34. Ilse, M., Tomczak, J. & Welling, M. Attention-based deep multiple instance learning. In International conference on machine learning, 2127–2136 (PMLR, 2018).

35. Lu, M. Y. et al. Data-efficient and weakly supervised computational pathology on whole-slide images. Nature biomedical engineering 5, 555–570 (2021).

36. Chen, R. J. et al. Scaling vision transformers to gigapixel images via hierarchical self-supervised learning. In Proceedings of the IEEE/CVF conference on computer vision and pattern recognition, 16144–16155 (2022).

37. Lipkova, J. et al. Deep learning-enabled assessment of cardiac allograft rejection from endomyocardial biopsies. Nature medicine 28, 575–582 (2022).

38. Shao, Z. et al. Transmil: Transformer based correlated multiple instance learning for whole slide image classification. Advances in neural information processing systems 34, 2136–2147 (2021).

39. Zhang, H. et al. Dtfd-mil: Double-tier feature distillation multiple instance learning for histopathology whole slide image classification. In Proceedings of the IEEE/CVF conference on computer vision and pattern recognition, 18802–18812 (2022).

40. Yan, R. et al. Shapley values-enabled progressive pseudo bag augmentation for whole-slide image classification. IEEE Transactions on Medical Imaging (2024).

41. Shao, D., et al. Do multiple instance learning models transfer? In Forty-second International Conference on Machine Learning (2025). URL https://openreview.net/forum?id=hfLqdquVt3.

42. Wang, X. et al. Transformer-based unsupervised contrastive learning for histopathological image classification. Medical image analysis 81, 102559 (2022).

43. Huang, Z., Bianchi, F., Yuksekgonul, M., Montine, T. J. & Zou, J. A visual–language foundation model for pathology image analysis using medical twitter. Nature medicine 29, 2307–2316 (2023).

44. Lu, M. Y. et al. A multimodal generative ai copilot for human pathology. Nature 634, 466–473 (2024).

45. Chen, R. J. et al. Towards a general-purpose foundation model for computational pathology. Nature Medicine 30, 850–862 (2024).

46. Xu, H. et al. A whole-slide foundation model for digital pathology from real-world data. Nature 630, 181–188 (2024).

47. Wang, X. et al. A pathology foundation model for cancer diagnosis and prognosis prediction. Nature 634, 970–978 (2024).

48. Ding, T. et al. Multimodal whole slide foundation model for pathology. arXiv preprint arXiv:2411.19666 (2024).

49. Xiang, J. et al. A vision–language foundation model for precision oncology. Nature 1–10 (2025).

50. Vaidya, A., et al. Molecular-driven foundation model for oncologic pathology. arXiv preprint arXiv:2501.16652 (2025).

51. Epstein, J. I. et al. A contemporary prostate cancer grading system: a validated alternative to the gleason score. European urology 69, 428–435 (2016).

52. Swanson, G. P., Trevathan, S., Hammonds, K. A., Speights, V. & Hermans, M. R. Gleason score evolution and the effect on prostate cancer outcomes. American journal of clinical pathology 155, 711–717 (2021).

53. Carroll, P. R. et al. NCCN guidelines insights: prostate cancer early detection, version 2.2016. Journal of the National Comprehensive Cancer Network 14, 509–519 (2016).

54. Schaeffer, E. M. et al. NCCN guidelines® insights: prostate cancer, version 3.2024: featured updates to the NCCN guidelines. Journal of the National Comprehensive Cancer Network 22, 140–150 (2024).

55. Jain, S. & Dhingra, S. Pathology of esophageal cancer and barrett’s esophagus. Annals of cardiothoracic surgery 6, 99 (2017).

56. Berry, M. F. Esophageal cancer: staging system and guidelines for staging and treatment. Journal of thoracic disease 6, S289 (2014).

57. Ajani, J. A. et al. Esophageal and esophagogastric junction cancers, version 2.2023, NCCN clinical practice guidelines in oncology. Journal of the National Comprehensive Cancer Network 21, 393–422 (2023).

58. Sun, X. & Xu, W. Fast implementation of DeLong’s algorithm for comparing the areas under correlated receiver operating characteristic curves. IEEE Signal Processing Letters 21, 1389–1393 (2014).

59. Sherstinsky, A. Fundamentals of recurrent neural network (RNN) and long short-term memory (LSTM) network. Physica D: Nonlinear Phenomena 404, 132306 (2020).

60. Odze, R. Diagnosis and grading of dysplasia in barrett’s oesophagus. Journal of clinical pathology 59, 1029–1038 (2006).

61. Fleiss, J. L. Measuring nominal scale agreement among many raters. Psychological bulletin 76, 378 (1971).

62. Vau, N. et al. Predicting biochemical recurrence after radical prostatectomy: the role of prognostic grade group and index tumor nodule. Human Pathology 93, 6–15 (2019).

63. Kim, G. et al. Holotomography. Nature Reviews Methods Primers 4, 51 (2024).

64. Park, J. et al. Revealing 3D microanatomical structures of unlabeled thick cancer tissues using holotomography and virtual H&E staining. Nature communications 16, 1–16 (2025).

65. Medina-Ramirez, I. E., Maćıas-Díaz, J. E., Masuoka-Ito, D. & Zapien, J. A. Holotomography and atomic force microscopy: a powerful combination to enhance cancer, microbiology and nanotoxicology research. Discover Nano 19, 64 (2024).

66. Teplov, A. et al. Development of standard operating procedure (SOP) of micro-computed tomography (micro-CT) in pathology. Diagnostic Pathology 5 (2019).

67. Rühli, F. J., Kuhn, G., Evison, R., Müller, R. & Schultz, M. Diagnostic value of micro-CT in comparison with histology in the qualitative assessment of historical human skull bone pathologies. American Journal of Physical Anthropology 133, 1099–1111 (2007).

68. Gao, G. et al. Comprehensive surface histology of fresh resection margins with rapid open-top light-sheet (otls) microscopy. IEEE Transactions on Biomedical Engineering 70, 2160–2171 (2023).

69. Chen, Y. et al. Rapid pathology of lumpectomy margins with open-top light-sheet (otls) microscopy. Biomedical optics express 10, 1257–1272 (2019).

70. Bishop, K. W. et al. Miniature line-scanned dual-axis confocal microscope for versatile clinical use. Biomedical Optics Express 14, 6048–6059 (2023).

71. Lu, M. Y. et al. A visual-language foundation model for computational pathology. Nature Medicine 30, 863–874 (2024).

72. Ertürk, A. Deep 3D histology powered by tissue clearing, omics and AI. Nature methods 21, 1153–1165 (2024).

73. Susaki, E. A. Unlocking the potential of large-scale 3d imaging with tissue clearing techniques. Microscopy (2024).

74. Wu, T. et al. Single-shot digital optical fluorescence phase conjugation through forward multiple-scattering samples. Science Advances 10, eadi1120 (2024).

75. Takanezawa, S., Saitou, T. & Imamura, T. Wide field light-sheet microscopy with lens-axicon controlled two-photon bessel beam illumination. Nature Communications 12, 2979 (2021).

76. Yu, Z., et al. Wavefront shaping: a versatile tool to conquer multiple scattering in multidisciplinary fields. The Innovation 3 (2022).

77. Krull, A., Buchholz, T.-O. & Jug, F. Noise2void-learning denoising from single noisy images. In Proceedings of the IEEE/CVF conference on computer vision and pattern recognition, 2129–2137 (2019).

78. Batson, J. & Royer, L. Noise2self: Blind denoising by self-supervision. In International conference on machine learning, 524–533 (PMLR, 2019).

79. Almagro-Pérez, C., et al. AI-driven 3D Spatial Transcriptomics. arXiv preprint arXiv:2502.17761 (2025).

80. Song, A. H., et al. Multimodal prototyping for cancer survival prediction. In Forty-first International Conference on Machine Learning (2024).

81. Jaume, G. et al. Transcriptomics-guided slide representation learning in computational pathology. In Proceedings of the IEEE/CVF Conference on Computer Vision and Pattern Recognition, 9632–9644 (2024).

82. Jaume, G. et al. Multistain pretraining for slide representation learning in pathology. In European Conference on Computer Vision, 19–37 (Springer, 2024).

83. Rao, V. M. et al. Multimodal generative AI for medical image interpretation. Nature 639, 888–896 (2025).

84. Barner, L. A. et al. Multiresolution nondestructive 3D pathology of whole lymph nodes for breast cancer staging. Journal of Biomedical Optics 27, 036501–036501 (2022).

85. Shao, J. & Tu, D. The jackknife and bootstrap (Springer Science & Business Media, 2012).

